# The Prion Protein Octarepeat Domain Forms Transient β-sheet Structures Upon Residue-Specific Cu(II) and Zn(II) Binding

**DOI:** 10.1101/2021.12.12.472308

**Authors:** Maciej Gielnik, Aneta Szymańska, Xiaolin Dong, Jüri Jarvet, Željko M. Svedružić, Astrid Gräslund, Maciej Kozak, Sebastian K. T. S. Wärmländer

## Abstract

Misfolding of the cellular prion protein (PrP^C^) is associated with the development of fatal neurodegenerative diseases called transmissible spongiform encephalopathies (TSEs). Metal ions appear to play a crucial role in the protein misfolding, and metal imbalance may be part of TSE pathologies. PrP^C^ is a combined Cu(II) and Zn(II) metal binding protein, where the main metal binding site is located in the octarepeat (OR) region. Here, we used biophysical methods to characterize Cu(II) and Zn(II) binding to the isolated OR region. Circular dichroism (CD) spectroscopy data suggest that the OR domain binds up to four Cu(II) ions or two Zn(II) ions. Upon metal binding, the OR region seems to adopt a transient antiparallel β-sheet hairpin structure. Fluorescence spectroscopy data indicates that under neutral conditions, the OR region can bind both Cu(II) and Zn(II) ions, whereas under acidic conditions it binds only Cu(II) ions. Molecular dynamics simulations suggest that binding of both metal ions to the OR region results in formation of β-hairpin structures. As formation of β-sheet structures is a first step towards amyloid formation, we propose that high concentrations of either Cu(II) or Zn(II) ions may have a pro-amyloid effect in TSEs.

## 1. Introduction

Transmissible spongiform encephalopathies (TSEs) are a group of neurodegenerative disorders initiated by misfolding of the cellular prion protein (PrP^C^)^1,2^. The human PrP^C^ is a 208 residues long protein expressed at high level in the central nervous system. It is composed of two structurally different regions: an unstructured N-terminal domain, and a globular and mostly α-helical C-terminal domain^3^ that attaches to the pre- and post-synaptic membranes via a GPI anchor^4,5^. For unknown reasons, and in a process similar to those for amyloid-forming peptides and proteins^6,7^, PrP^C^ can undergo a structural transition into an insoluble, aggregated form with high β-sheet content called PrP^Sc^. It has been suggested that PrP is most toxic when forming soluble oligomers, i.e. intermediate species during PrP^Sc^ formation, which can accumulate in brain tissue and cause neurodegeneration^8^. Even though human TSEs are very rare and only affect one person per million^9^, they share many similarities with the pathologies characterized by proteins aggregating into amyloid states, and multiple evidence indicates that the prion and amyloid diseases all belong to a large family of protein aggregation diseases^10–13^. Examples include tauopathies (tau protein)^14^, Alzheimer’s disease (Aβ peptide)^15^, Parkinson’s disease (α-synuclein protein)^16^, and amyotrophic lateral sclerosis/ALS (TDP-43 protein)^17,18^. Thus, it has recently been proposed that beside TSEs, PrP can be involved also in the development of other neurodegenerative diseases, such as Alzheimer’s disease^19^.

The native function of PrP^C^ is still elusive. The PrP^C^ protein is encoded by the *PRNP* gene, and most *PRNP* knockout animals (i.e., without the *PRNP* gene) show normal development and behavior, although some individuals show deviation in neuronal signal transduction and locomotion^20^. Interestingly, all *PRNP* knockout mice are immune to PrP^Sc^ inoculation, which supports the theory of template-driven autocatalytic conversion of PrP^C^ to PrP^Sc2,11^. Among the many functions attributed to PrP^C^, i.e. cell signaling, antioxidation, and myelination^8^, phylogenetic analysis indicates that PrP^C^ is evolutionary linked to the Zrt- and Irt-like Protein (ZIP) family of divalent metal transporters^21^. This suggests that one role for PrP^C^ might be in metal ion homeostasis, and metal imbalance has been suggested to be part of the pathology of prion diseases^22–24^

The PrP^C^ protein binds up to six different types of divalent metal ions, including Cu(II), Zn(II), Ni(II), and Mn(II), by two distinct domains with different metal ion affinities^26–28^. The octarepeat (OR) region is located in the N-terminal domain where it spans residues 60-91 (Fig. 1). It contains four tandem PHGGGWGQ repeats, and binds Cu(II), Zn(II) and Ni(II) ions with strong affinity^29^. The so-called “non-octarepeat region” spans residues 92-111, and binds Cu(II) ions with weaker affinity, where H96 and H111 are likely binding ligands^30,31^. The capacity of the OR region to bind Cu(II) ions has been intensively studied during the last twenty years. Cu(II) is an important neurotransmitter and the third most common transition metal in the brain^32^. The reported Cu(II) concentration in the synaptic cleft during neuron depolarization ranges from 3 μM^33^ to 250 μM^34^, which suggests that the dissociation constant (K_d_) for the OR·Cu(II) complex is at least of this order of magnitude. The OR region has been reported to bind up to four Cu(II) ions^35–37^, where the first ion is bound with the highest affinity (around 0.1 nM), and the three other display intermediate affinities around 10 μM^38^. Many possible biological functions have been attributed to the Cu(II)-binding capacity of PrP^C^, including superoxide dismutase activity, transmembrane copper transport, copper buffering, and neuronal protection ^39–42^.

**Figure 1.**
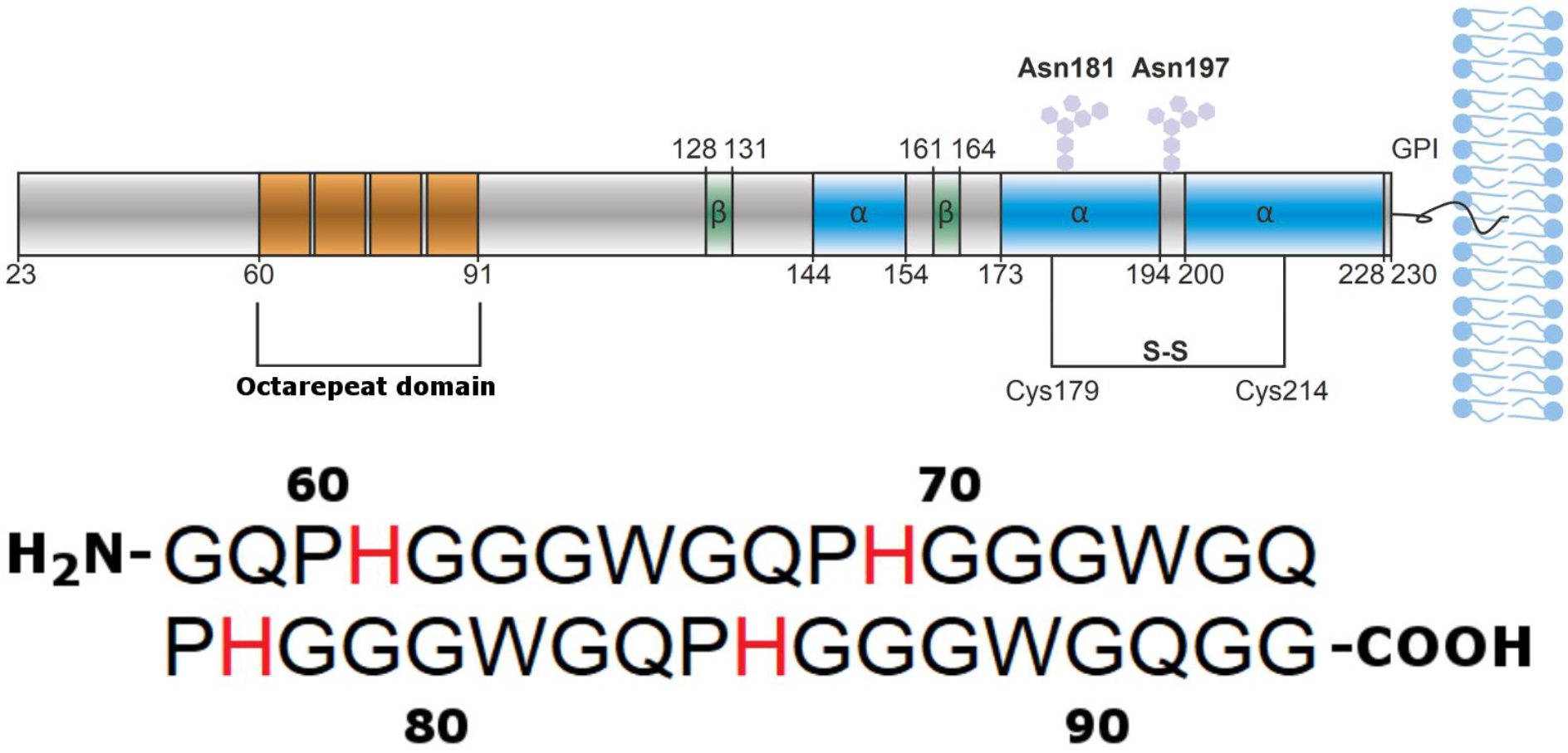
Top: Sequence of the human prion protein, with the octarepeat region marked as orange, β-sheets marked as green, and α-helices marked as blue. Image is from Gielnik et al.^25^ under CC BY 4.0. Bottom: The octarepeat (OR) region studied in this paper comprises residues 58 – 93 of the prion protein, i.e. PrP^C^(58-93). At neutral pH it has no charged residues. Possible metal-binding aromatic histidine residues are shown in red.

Another important metal ion for PrP^C^ neurobiology is Zn(II). Zinc is the second-most (after iron) abundant metal ion in the human body^43^. Upon neuronal stimulation, the transient concentration of Zn(II) ions in the synaptic vesicle can reach values around 300 μM^44^. Such Zn(II) concentrations stimulate PrP^C^ endocytosis into human neuroblastoma cells ^45^, and PrP^C^ has been shown to enhance Zn(II) transport^46^. An early study reported a K_d_ of 200 μM for the PrP^C^·Zn(II) complex^47^, but more recent ITC studies by the same researchers suggest a K_d_ of 17 μM, together with NMR experiments implying 1:1 stoichiometry^48^. The proposed PrP^C^ functions related to Zn(II) binding are similar to those proposed for Cu(II) binding, e.g. metal ion buffering and transport^24,49^.

We have recently shown that the isolated OR region (i.e., an OR peptide) upon interaction with Zn(II) ions forms fibrillar cross-β structures that bind thioflavin T and Congo Red, and which possess all the characteristic features of amyloid material^50^. This indicates that metal ions can directly induce a transition of PrP^C^ into the amyloid state.

Here, we use circular dichroism (CD) and fluorescence spectroscopy, combined with molecular dynamics simulations, to estimate apparent K_d_ values for the OR·Cu(II) and OR·Zn(II) complexes, and to characterize the initial structural changes of the isolated OR region, i.e. PrP^C^(58-93) (Fig. 1), after exposure to Cu(II) and Zn(II) ions.

## 2. Materials and Methods

### 2.1. Peptide synthesis and purification

The OR peptide, PrP(58-93), was obtained using solid peptide synthesis methodology. TentaGel R RAM resin (loading capacity of 0.18 mmol/g; Rapp Polymere, Germany) was used as a matrix. The synthesis was performed using a standard Fmoc/tBu amino acid chemistry on Microwave Liberty Blue synthesizer (CEM) peptide synthesizer. The crude peptide was cleaved from the solid support using a cleavage cocktail consisting of 88% trifluoroacetic acid (TFA), 5% water, 5% phenol and 2% triisopropylsilane (v/v/m/v) under neutral (argon) atmosphere, and protected from direct exposure to light. After precipitation with diethyl ether, the crude product was dissolved in water and lyophilized.

Peptide purification was carried out by reversed-phase high-performance liquid chromatography (RP-HPLC), using a Luna C8(2) AXIA Pack column (250 x 21.2 mm, 5 μm, 100 Å; Phenomenex, USA). A linear gradient of acetonitrile in 0.1% aqueous TFA was applied as a mobile phase. The purity of the obtained fractions was evaluated by analytical UHPLC, using a Kinetex C8 column (100 x 2.1 mm, 2.6 μm, 100 Å; Phenomenex, USA) operated by the NEXERA-i chromatography system (Shimadzu, Japan) in a 15 min linear gradient of 5-100% B (where B is 80% acetonitrile in 0.1% aqueous TFA). UV absorption was monitored at λ = 223 nm. Fractions with a purity higher than 99 % were pooled together for further analyses. The molecular weight of the final peptide sample was confirmed by mass spectrometry, using an ESI-IT-TOF-LC-MS system (Shimadzu, Japan) with a C12 Jupiter Proteo column (150 × 2 mm, 4μm, 90 Å; Phenomenex, USA).

### 2.2. Circular dichroism spectroscopy

Circular dichroism (CD) spectra of the OR peptide were recorded on a Chirascan CD (Applied Photophysics, UK) spectropolarimeter. Thermal unfolding experiments were performed for 20 μM OR peptide in 10 mM sodium phosphate buffer, pH 7.0, in a cuvette with 4 mm pathlength, in the range from 5 °C to 65 °C with 5 °C intervals. Spectra were recorded from 190 nm to 250 nm, with 0.5 nm steps and a time-per-point of 4 s. The content of PPII helix was estimated from the CD intensity expressed in mean residue ellipticity (a concentration-independent unit) at the local maximum at 225 nm of the CD spectra, i.e. θ_max_, using the equation published by Kelly et al.^51^, i.e., eq. 1:

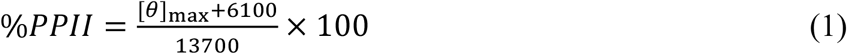

Titrations with metal ions were conducted for 5 μM OR peptide dissolved either in pure MilliQ water or in 10 mM sodium phosphate buffer, pH 7.5. Using 1 cm path-length quartz cuvettes with gentle magnetic stirring at 25°C, the OR peptide (volume 2.5 ml) was titrated with small amounts of stock solutions of CuCl_2_ or ZnCl_2_ (100 μM,500 μM, 1.25 μM or 5 μM stock concentrations) directly in the cuvette. All spectra were collected from 200 to 260 nm with sampling points every 0.5 nm, a time-per-point of 4 s, and 2 nm bandwidth. The final spectra were baseline-corrected and smoothed with a Savitzky-Golay filter. Data with single visible transitions were fitted to the transformed Hill equation, i.e., eq. 2:

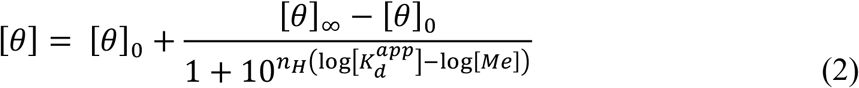

Here, [θ]0 is the signal intensity before the transition, [θ]∞is the signal intensity at the end of the transition, n_H_ is the Hill coefficient, [K_d_^app^] is the apparent dissociation constant, and [Me] is the metal ion concentration. When two transitions were observed, and the signal was monotonically increasing or decreasing, the data was fitted as a sum of two transformed Hill equations, i.e., eq. 3:

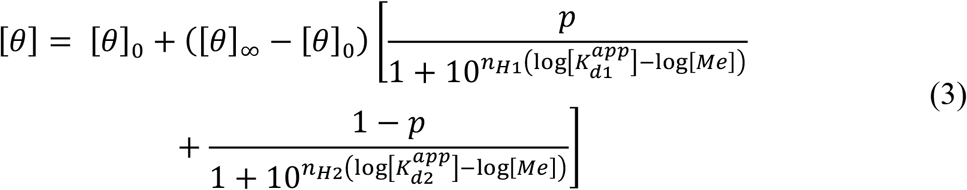

Here, [θ]_0_ is the signal intensity before the transition, [θ]_∞_ is the signal intensity at saturation, [K_d1_^app^] and [K_d2_^app^] are the apparent dissociation constants for the first and second binding sites, n_H1_ and n_H2_ are the Hill coefficients for the first and second binding sites, [Me] is the metal ion concentration, and p and 1-p are the relative signal intensities for the first and second binding sites. When the two transitions were observed with a local maximum or minimum in the signal, data was fitted as the sum of one transformed and one reverse-transformed Hill equation, i.e., eq. 4:

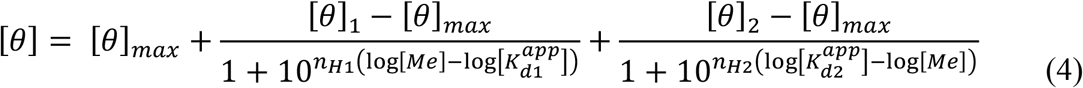

Here, [θ]_max_ is the signal intensity at the extreme point, [θ]_1_ is the signal intensity before the transition, [θ]_2_ is the signal intensity after the transition, and n_H1_, n_H2_, [K_d1_^app^], [K_d2_^app^], and [Me] have the same meaning as in eq. (3).

### 2.3. Fluorescence Spectroscopy

Fluorescence spectra of the OR peptide were recorded with a Cary Eclipse (Varian, USA) fluorometer equipped with a Peltier multicell holder, using quartz cuvettes with 1 cm path length. 5 μM of OR peptide was titrated with stock solutions of CuCl_2_ and ZnCl_2_, similar to the CD titrations (above), and using the following different buffers: (i)10 mM sodium phosphate buffer, pH 7.5, (ii) 10 mM MES (2-morpholinoethanesulfonic acid) buffer, pH 5.5, (iii) 10 mM MES buffer, pH 7.5. Some titrations with CuCl_2_ were conducted in the presence of 1 mM of the reducing agent TCEP (tris(2-carboxyethyl)phosphine), to investigate the effect of reduced Cu(I) ions. The samples were excited at 285 nm, and emission spectra were recorded in the 300 - 500 nm range, with a 1 nm data interval and a scan speed of 600 nm/min. The excitation and emission bandwidths were respectively 10 nm and 5 nm, and the experiments were performed at 25 °C under quiescent conditions (i.e., no stirring). The fluorescence intensity at 255 nm was plotted versus metal ion concentration, and the resulting data curves were fitted with the same equations as those used for CD data (i.e., eq. 2).

### 2.4. Molecular dynamics simulations

Molecular dynamics simulations were performed in GROMACS 2019.2^52^ using the OPLS-AA^53^ force field. The systems were solvated with the TIP4P^54^ water model and restrained using Van der Waals radii^55^. The LINCS algorithm^56^ was used to restrain all covalent bonds in the peptide, and the SETTLE algorithm^57^ was used to restrain all water molecules. In the initial model one or two nonbonded dummy Cu(II) or Zn(II) ion models^58^ were placed in close proximity to N^ε2^ atoms of histidine side chains. The N^ε2^ atoms of the four histidine residues were either protonated or uncharged, to study the interactions under different pH conditions (for short peptides histidine protonation/deprotonation correspond to pH < 6.8 or pH > 6.8, respectively)^59^. The systems were neutralized with Cl^-^ ions, and the energy was minimized using steepest-descent energy minimization over 5000 steps. The temperature was equilibrated at 300 K, in an NVT ensemble over 0.5 ns with 1 fs time steps, using the modified Berendsen^60^ thermostat. The pressure was equilibrated at 1 bar, in NPT ensemble over 0.5 ns with 1 fs time steps, using the Parrinello-Rahman^61^ barostat. For long-range electrostatic interactions we applied PME^62^ with 1.2 nm cutoff, and the same cutoff was used for Van der Waal forces. The MD production runs were performed in an NVT ensemble with 2 fs time step. The trajectories for the peptide with uncharged histidine residues were produced over 100 ns, while trajectories for the peptide with protonated histidine residues were produced over 10 ns. The results were analyzed in VMD^63^ and visualized in the PyMOL Molecular Graphics System, Version 2.3.4 (Schrödinger, LLC). The secondary structure was assigned using the PROSS software^64^, which estimates secondary structures as α-helix, β-strand, β-turn, and PPII, and classifies all other structures as “coil”. The principal component analysis was performed in the Bio3D R package^65^.

### 2.5 Calculation of pKa values

Final peptide conformations were extracted as frames from MD simulations by storing the structures as PDB files, which then were used as input for the two best known protocols for pKa calculations for protein structures. Thus, pKa values for the OR peptide were calculated using both the PropKa 2.0 software^66^ with the PARSE^67^ force field, and the DelPhiPKa software^68^ with the AMBER force field^69^. The pKa values were calculated at physiological pH and ionic strength at 37 °C, following the instructions for each program. Two independent protocols were used for evaluation of calculation accuracy.

## 3. Results

### 3.1. The octarepeat (OR) region exhibits a mixture of random coil and PPII helix

The initial low-temperature (5 °C) CD spectrum of 20 μM *apo*-OR peptide showed one positive band at 225 nm, together with two negative bands: a weak one at 238 nm, and a strong one at 199 nm (Fig. 2A, dark green spectrum). The minimum around 199 nm is consistent with a random coil structure, but this conformation should not give rise to positive CD bands, such as the one at 225 nm. Other researchers have previously suggested that the OR peptide in aqueous solution adopts a PPII left-handed extended helix^70,71,72^, or exhibits a mixture of random-coil and β-turn structures^35,73,74^. To clarify the secondary structure of the OR peptide, we recorded CD spectra at different temperatures to monitor the thermal unfolding of the peptide (Figs. 2A and 2B).

**Figure 2.**
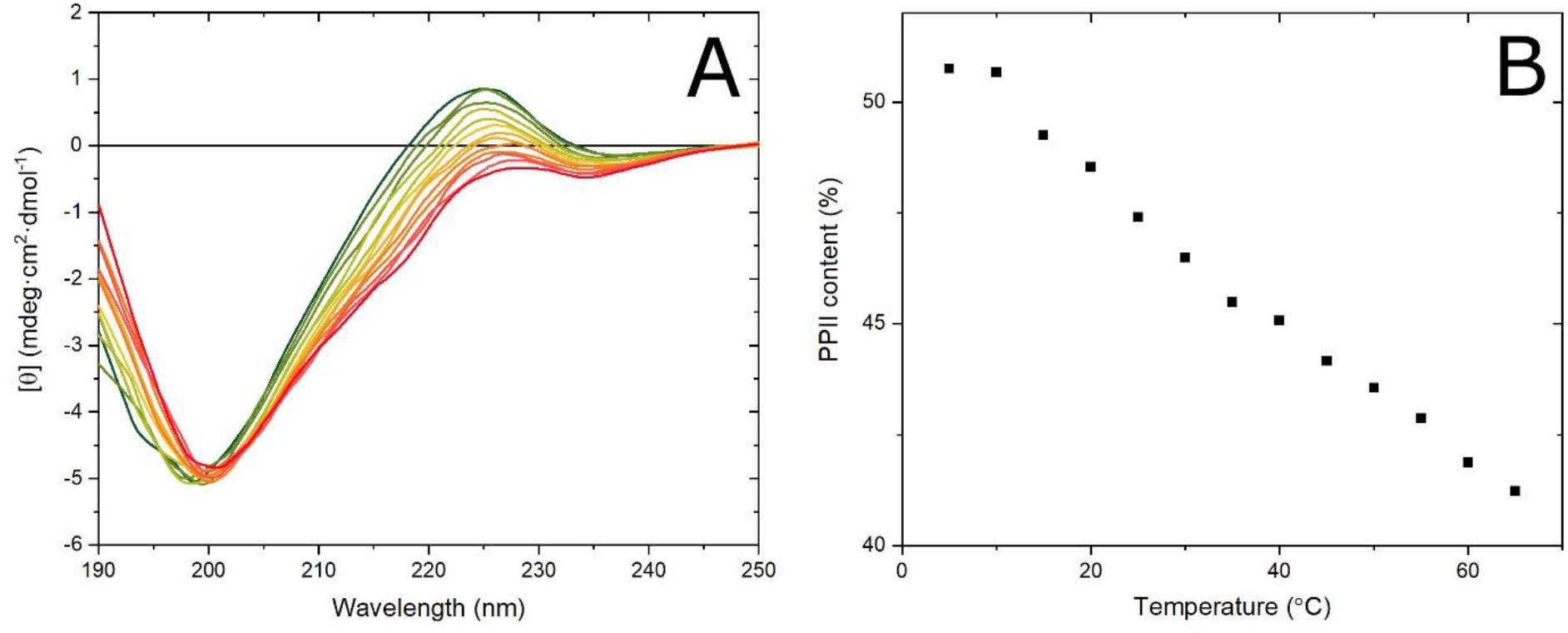
(**A**) CD spectra for thermal unfolding of 20 μM OR peptide from 5 °C (dark green line) to 65 °C (red bottom line) at 5 °C intervals. The isodichroic point at 204 nm, for spectra from 20 °C to 65 °C, suggests a PPII helix to random coil transition. All CD spectra were recorded in 10 mM phosphate buffer, pH 7.0. (**B**) Estimated content of PPII helix for all recorded temperatures, calculated from eq. 1 and CD intensities at 225 nm.

The intensity of the CD spectra at 225 nm gradually decreased when the temperature increased from 5 °C to 20 °C, but the spectral quality did not allow for any detailed interpretation of the spectral shape (Fig. 2A). For the spectra between 20 °C and 65 °C, however, an isodichroic point appeared at 204 nm (Fig. 2A). Together with the gradual decrease of the 225 nm band, this indicates a structural transition from PPII helix to random coil conformation^75^. We therefore calculated the PPII content in the OR peptide as a function of temperature (Fig. 2B), using the CD signal intensity at 225 nm and eq. 1.^51^ The PPII content was highest at 5 °C, i.e. around 51%, and then gradually decreased to around 41% at 65 °C (Fig. 2B). Interestingly, in the temperature range of the isodichroic point, i.e. 20 °C to 65 °C, the amount of PPII helix decreased linearly as a function of temperature. Overall, these CD results indicate that the secondary structure of the OR peptide at physiological temperature is a mixture of random coil and PPII helix, with a significant amount - more than 45% - of PPII structure.

### 3.2. Cu(II) and Zn(II) binding to the OR peptide both induce formation of antiparallel β-sheet structure

Previous studies suggest that Cu(II) binding to the OR peptide in pure water induces certain changes in the peptide’s secondary structure, involving formation of β-turns or structured loops around the metal ions^35^. Our initial titrations of 5 μM OR peptide with CuCl_2_ in water (pH adjusted to ~7.5 with NaOH and controlled by a pH meter) showed a gradual decrease in the CD signal that can be associated with peptide precipitation (Figure S1A). The final spectrum had a weak single minimum at 220 nm and a maximum at 208 nm, which may suggest formation of β-sheets. As the direct titrations with CuCl_2_ in water appeared to induce severe aggregation, a different approach was tried. The 5 μM peptide solution was acidified with small amounts of acetic acid to pH ~4.0, then 20 μM of CuCl_2_ was added, and the pH of the solution was slowly increased to ~7.5 with NaOH. This approach allowed us to acquire a CD spectrum with a shape that previously has been described in the literature as typical for the OR:Cu(II) complex, and interpreted as formation of β-turns or structured loops^35,72,73^ (Fig. 3A, red spectrum).

**Figure 3.**
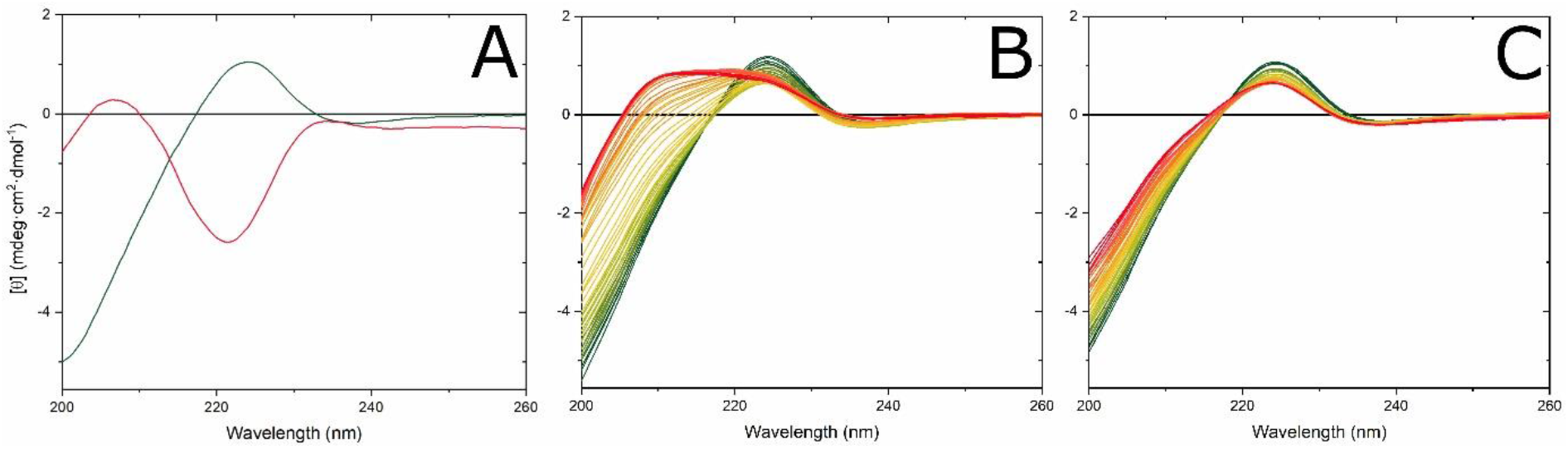
CD spectra for Cu(II) and Zn(II) binding to the OR peptide at 25 °C. (**A**) 5 μM OR peptide in water at pH 7.5 (no buffer) before (green line) and after (red line) addition of 20 μM CuCl_2_. CuCl_2_ was added at a pH of 4.0, and then the pH was adjusted to 7.5 with small amounts of NaOH. The shape of the red CD spectrum shows features reported in the literature as typical for a Cu(II)-OR complex. (**B**) Titration of 5 μM OR peptide in 10 mM phosphate buffer, pH 7.5, with CuCl_2_ from 0 μM (green) to 40 μM (red). (**C**) Titration of 5 μM OR peptide in 10 mM phosphate buffer, pH 7.5, with ZnCl_2_ from 0 μM (green) to 40 μM (red).

To investigate the binding of Cu(II) ions to the OR peptide in a more controlled environment, we titrated CuCl_2_ to 5 μM OR peptide in 10 mM phosphate buffer, pH 7.5 at 25 °C (Fig. 3B). The initial CD spectrum then began to lose intensity as 224 nm, and a new band appeared at 208 nm. Careful analysis of these CD spectra revealed three distinct spectral transitions, likely corresponding to transitions in the peptide’s secondary structure. The first transition was present from 0 μM up to 5 μM of CuCl_2_, corresponding to 1:1 Cu(II):OR peptide ratio. During this process the CD intensity at 224 nm decreased, and a new weak band appeared at 208 nm (Fig. 3B, green to yellow spectra). The isodichroic point at 217 nm was clearly visible, which could suggest a PPII to β-turn structural transition^76^.

The second spectral transition appeared at CuCl_2_ concentrations from 5 μM to 10 μM, corresponding to 2:1 Cu(II):OR peptide ratio. During this process the CD intensity at 208 nm strongly increased, while the CD band at 224 nm showed a small intensity increase (Fig. 3B, yellow to orange spectra). The absence of an observed isodichroic point excluded the possibility of a two-state transition and suggests formation of a new CD band. The difference spectrum for this transition (Fig. S2B, red line) resembled the CD spectrum for relaxed antiparallel β-sheets^77^. On the other hand, the difference spectrum could also be an inverted CD spectrum of a random coil^76^, which would imply a loss in random coil secondary structure. However, the CD spectrum of a random coil secondary structure should have a minimum below 200 nm, but in our case the extremum is at 205 nm (Fig. S2B, red line). We therefore suggest that the second spectral transition involves formation of a relaxed antiparallel β-sheet secondary structure.

The third transition appeared for CuCl_2_ concentrations from 10 μM to 40 μM (Fig. 3B, orange to red spectra). During this process, the newly formed band at 208 nm reached a maximum and maintained constant intensity, where the band at 224 nm began to lose intensity. No isodichroic point was observed, similarly to the second transition, which again suggests formation of a new spectral band. The difference spectrum (Fig. S2B, blue line) showed features similar to those observed in the second spectral transition, again suggesting formation of relaxed antiparallel β-sheets^77^. However, as the changes in the CD spectral intensity were smaller for the third transition then for the second transition, less of the new structure appears to have formed.

The final CD spectrum for the OR·Cu(II) complex in the phosphate buffer has two maxima at 224 nm and 208 nm (Fig. 3B, red spectrum), whereas the CD spectrum for the OR·Cu(II) complex in pure water has a maximum at 208 nm and a minimum at 220 nm (Fig. 3A, red spectrum). Thus, the presence of the buffer may influence the structural transitions in the OR peptide during Cu(II) binding. The difference spectrum for the OR·Cu(II) complex in water has a maximum at 202 nm and a minimum at 222 nm (Fig. S2A), and thus resembles the CD spectrum of a left-hand twisted antiparallel β-sheet, which is supposed to have a maximum at ~203 nm and a minimum at ~226 nm^77^. This suggests that both in water and phosphate buffer, Cu(II) binding to the OR peptide results in formation of antiparallel β-sheet structures, however possibly with different geometries, i.e., relaxed antiparallel β-sheets vs. left-hand twisted antiparallel β-sheets.

Interestingly, titrating 5 μM OR peptide in the phosphate buffer with CuCl_2_ up to 40 μM, in 5 μM intervals, produced different CD spectra (Fig. S1B), than the CuCl_2_ titrations with very small steps (Fig. 3B). The final CD spectrum in the small-step titration had a maximum at 224 nm, and formation of a new band at 208 nm was visible (Fig. 3B, red spectrum). For the titrations with larger steps, i.e. 5 μM intervals, the isodichroic point at 217 nm was absent, the new band formed at 208 nm had lower intensity, and the band at 224 nm was not affected by the Cu(II) ions at all. The difference spectrum between these two titrations (Fig. S1B, blue line) shares similarities with the difference spectra for titrations in the phosphate buffer with small steps (Fig. S2B, red line, blue line), thus suggesting formation of antiparallel relaxed β-sheets. We therefore suggest that during the large-step titrations, Cu(II) ions may have been bound in a rather chaotic way, possibly favoring intermolecular binding conformations that would induce peptide aggregation.

The titrations of the OR peptide with Cu(II) ions using small steps (Fig. 3B) resembled a steady-state approximation, wherefore we analyzed these data in a more quantitative manner. Plotting the mean residue ellipticity at 208 nm and 224 nm versus CuCl_2_ concentration shows all three spectral transitions in the OR peptide (Figs. 4A and 4B). The data at 208 nm fitted with eq. 3 yields apparent dissociation constants of 0.52 μM for the first transition, and 5.57 μM for the second transition. The change in CD intensity at 224 nm for the first transition fitted with eq. 2 produced an K_d1_^app^ of 0.24 μM. The second and third transitions, visible at 224 nm and fitted to eq. 4, produced the values K_d2_^app^ = 7.84 μM and K_d3_^app^ = 18.0 μM (Table 1). Thus, both the 208 nm and 224 nm CD data produced affinity values that were in the sub-micromolar range for the first transition and in the low micromolar range for the second transition. Moreover, the 208 nm and 224 nm CD data produced also similar Hill coefficients. The Hill coefficient for the first transition (PPII helix to β-turn) oscillates around 1, suggesting a single binding site. For the second and third transitions, the Hill coefficients are larger than 1, suggesting multiple binding sites (Table 1).

**Figure 4.**
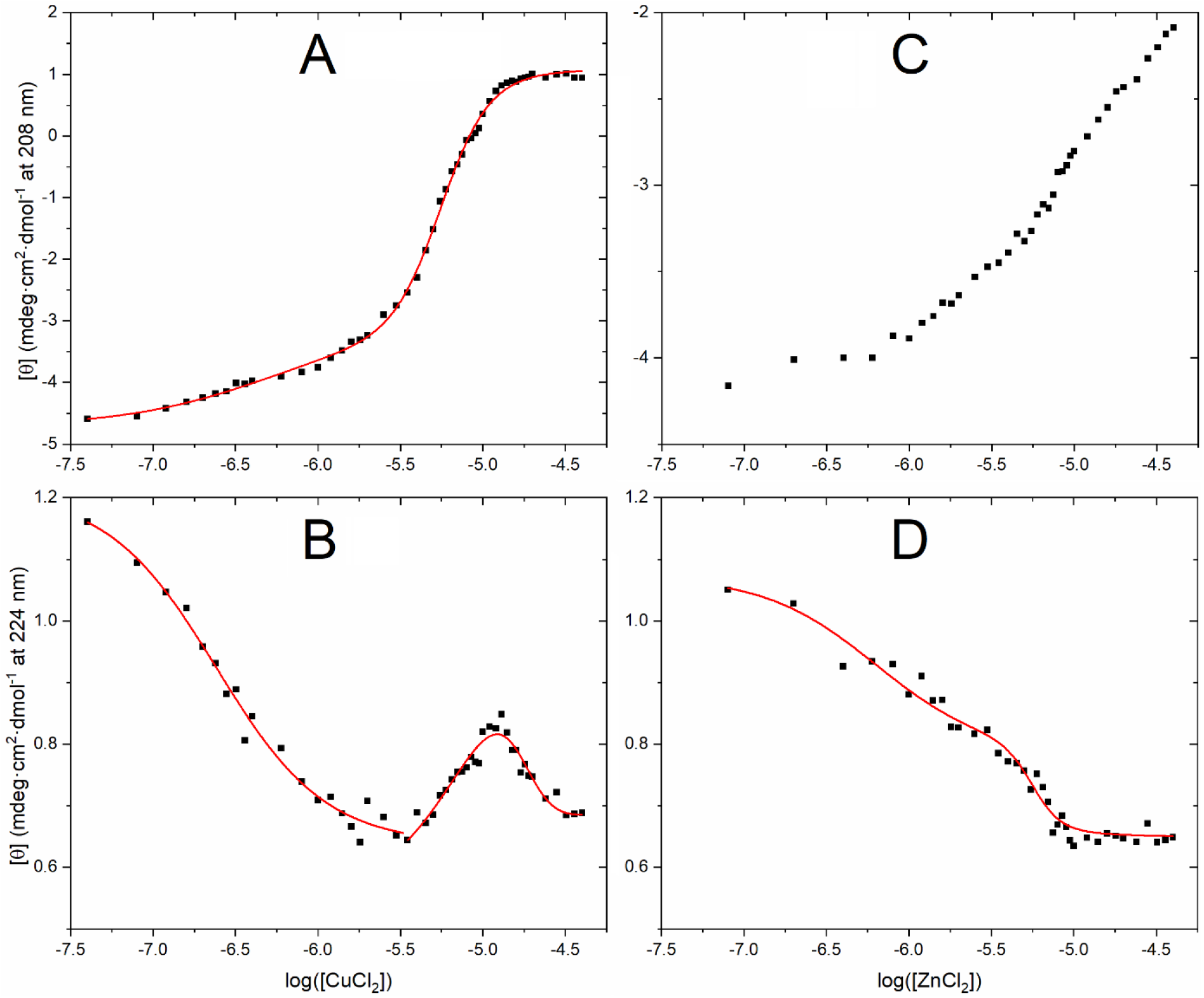
Changes in CD intensity [θ] for 5 μM OR peptide at 25 °C in 10 mM phosphate buffer, pH 7.5, when titrated with metal ions as shown in Fig. 3. (**A**) 208 nm for Cu(II) ions (Fig. 3B); (**B**) 224 nm for Cu(II) ions (Fig. 3B); (**C**) 208 nm for Zn(II) ions (Fig. 3C); (**D**) 224 nm for Zn(II) ions (Fig. 3C).

**Table 1.** Apparent dissociation constants (K_d_^app^) and Hill coefficients (n_H_) for the OR-metal ion complex, based on fluorescence quenching and circular dichroism (CD) experiments.

| Metal ion | Method | Buffer | $K_{d1}^{app}$ [μM] | $n_{H1}$ | $K_{d2}^{app}$ [μM] | $n_{H2}$ | $K_{d3}^{app}$ [μM] | $n_{H3}$ |
| --- | --- | --- | --- | --- | --- | --- | --- | --- |
| Cu(II) | CD at 208 nm | 10 mM NaH<sub>2</sub>PO<sub>4</sub>, pH 7.5 | 0.52 ± 0.31 | 0.97 ± 0.48 | 5.57 ± 0.11 | 2.97 ± 0.25 | | |
| Cu(II) | CD at 224 nm | 10 mM NaH<sub>2</sub>PO<sub>4</sub>, pH 7.5 | 0.24 ± 0.03 | 1.31 ± 0.20 | 7.84 ± 0.25 | 1.81 ± 0.14 | 18.0 ± 0.7 | 4.50 ± 0.41 |
| Zn(II) | CD at 224 nm | 10 mM NaH<sub>2</sub>PO<sub>4</sub>, pH 7.5 | 0.64 ± 0.17 | 1.28 ± 0.26 | 5.61 ± 0.37 | 5.11 ± 2.02 | | |
| Cu(II) | Fluorescence at 355 nm | 10 mM NaH<sub>2</sub>PO<sub>4</sub>, pH 7.5 | 4.29 ± 0.06 | 1.5 ± 0.03 | | | | |
| Cu(II) | Fluorescence at 355 nm | 10 mM MES, pH 7.5 | 4.5 ± 0.1 | 1.61 ± 0.05 | | | | |
| Cu(II) | Fluorescence at 355 nm | 10 mM MES, pH 5.5 | 7.4 ± 0.1 | 1.21 ± 0.02 | | | | |
| Cu(II) | Fluorescence at 355 nm | 10 mM MES, pH 7.5 | 81 ± 31 | 0.61 ± 0.03 | | | | |

To investigate the Zn(II) binding to the OR peptide we titrated 5 μM OR peptide with ZnCl_2_ in the phosphate buffer, pH 7.5, at 25 °C (Fig. 3C). During the titrations with Zn(II) ions, the CD spectra of the OR peptide gradually lost intensity at 224 nm, and a weak new band appeared at 208 nm. Plotting the CD signal intensities at 208 nm and 224 nm versus the ZnCl_2_ concentration clearly showed three spectral transitions (Figs. 4C and 4D). The data at 224 nm showed two transitions, and fitting to eq. 3 produced apparent dissociation constants of 0.64 μM and 5.61 μM for the first and second transitions (Table 1). The CD intensity at 208 nm did not show a saturation or inflection point in the studied concentration range of ZnCl_2_, and was therefore not used for calculating binding affinity values. We may however speculate that this signal could have an inflection point above 40 μM, corresponding to rather weak binding.

The first transition appeared from 0 μM to 5 μM of ZnCl_2_, (Fig. 3C, green to yellow spectra). During this process the CD intensity at 224 nm decreased, and a new weak band at 208 nm appeared. A clear isodichroic point at 217 nm was visible, suggesting a PPII to β-turn structural transition^76^.

The second transition was visible from 5 μM to 10 μM of ZnCl_2_ (Fig. 3C, yellow to orange). During this process the CD signal intensity at 224 nm and 208 nm decreased (Fig. 4C and 4D), while the isodichroic point gradually shifted from 217 nm to 218.5 nm. The new isodichroic point suggests a two-state structural transition, different from the PPII to β-turn transition. The difference spectrum for this transition had one minimum at 226 nm, and at least two maxima, one at 208 nm and a second below 200 nm (Fig. S2C, red line). The maximum at 208 nm and the minimum at 226 nm might correspond to a left-handed twisted antiparallel β-sheet, suggesting that such a structure could be induced by higher concentrations of Zn(II) ions.

The third transition was observed for ZnCl_2_ concentrations above 10 μM (Fig. 3C, orange to red). In this spectral transition no clear isodichroic point was present, and the band at 208 nm gradually lost intensity, while the signal at 224 nm remained constant (Figs. 4C and 4D). The difference CD spectrum between the start and end points of this transition (Fig. S2C, blue line) was similar to the difference spectrum for the OR peptide titrated with Cu(II) ions (Fig. S2B, blue line), suggesting that this transition also involves formation of antiparallel β-sheets.

Interestingly, the final CD spectrum for the Zn(II) titration, i.e. the OR peptide with 40 μM ZnCl_2_, is nearly identical to the CD spectrum for the OR peptide with 4 μM CuCl_2_ (Figs. 3B and 3C). Characteristic features are here decreased signal intensity at 224 nm and formation of a new band at 208 nm. But for the Cu(II) titration, this is just an intermediate step – the final step with 40 μM CuCl_2_ shows a very strong increase of the 208 nm band, which is something that Zn(II) ions apparently are not able to induce.

### 3.3. Protonation of His residues decreases the OR affinity for Cu(II) and Zn(II) ions

Tryptophan residues have previously been shown to indirectly coordinate Cu(II) ions in a HGGGW sequence, corresponding to the core of a single isolated octarepeat sequence^36^, and quenching of tryptophan fluorescence has been successfully applied to calculate the affinity between the octarepeat region and metal ions^29,38,78,79^. Thus, we here used fluorescence spectroscopy measurements of the intrinsic tryptophan fluorescence to further investigate binding of Cu(II) and Zn(II) ions to the OR peptide. Cu(II) is a paramagnetic ion, and can therefore strongly quench nearby fluorophores. Zn(II) is not paramagnetic, and any changes in tryptophan fluorescence should therefore originate only from zinc-induced changes in the peptide structure.

Cu(II) ions were titrated to 5 μM OR peptide at 25 °C in three sets of buffers: 10 mM phosphate buffer, pH 7.5; 10 mM MES buffer, pH 7.5 and 10 mM MES buffer, pH 5.5. As we believe that the OR peptide coordinates Cu(II) ions via histidine sidechains^36,37,80^, we expect the binding to be weaker at pH 5.5, where the His residues will become protonated^59^. In all three buffers Cu(II) ions were found to quench the intrinsic tryptophan fluorescence, clearly demonstrating that Cu(II) ions bind to the OR peptide in all the studied conditions. In all cases the maximum fluorescence intensity was at 355 nm, and this maximum did not change its position during the titrations. In buffers at pH 7.5, the Cu(II) ions quenched the tryptophan fluorescence to 30% of the initial intensity (Figs. 5A and 5B), while in 10 mM MES buffer at pH 5.5, the tryptophan quenching was reduced to 50% of the initial fluorescence intensity (Fig. 5C). To derive apparent K_d_ values, we plotted the fluorescence intensity at 355 nm as a function of the CuCl_2_ concentration and fitted the data to eq. 2 (Figs. 5A, 5B, and 5C: insets). The calculated K_d_^App^ values were similar in the phosphate buffer and in the MES buffer at neutral pH, i.e. respectively 4.3 μM and 4.5 μM (Table 1). At acidic conditions the calculated K_d_^App^ was somewhat higher, i.e. 7.5 μM, suggesting a slightly weaker binding. The K_d_^App^ values derived with fluorescence spectroscopy are roughly one order of magnitude higher than those derived with CD spectroscopy, which is something that we attribute to differences in the two methods, although it should be pointed out that different buffers were used (Table 1). MES is a “Good” buffer with minimal metal binding^81^, but it is not suitable for CD measurements. Phosphate buffer is compatible with most spectroscopic techniques, and biologically relevant, but the phosphate ions may have some interactions with metal ions.

**Figure 5.**
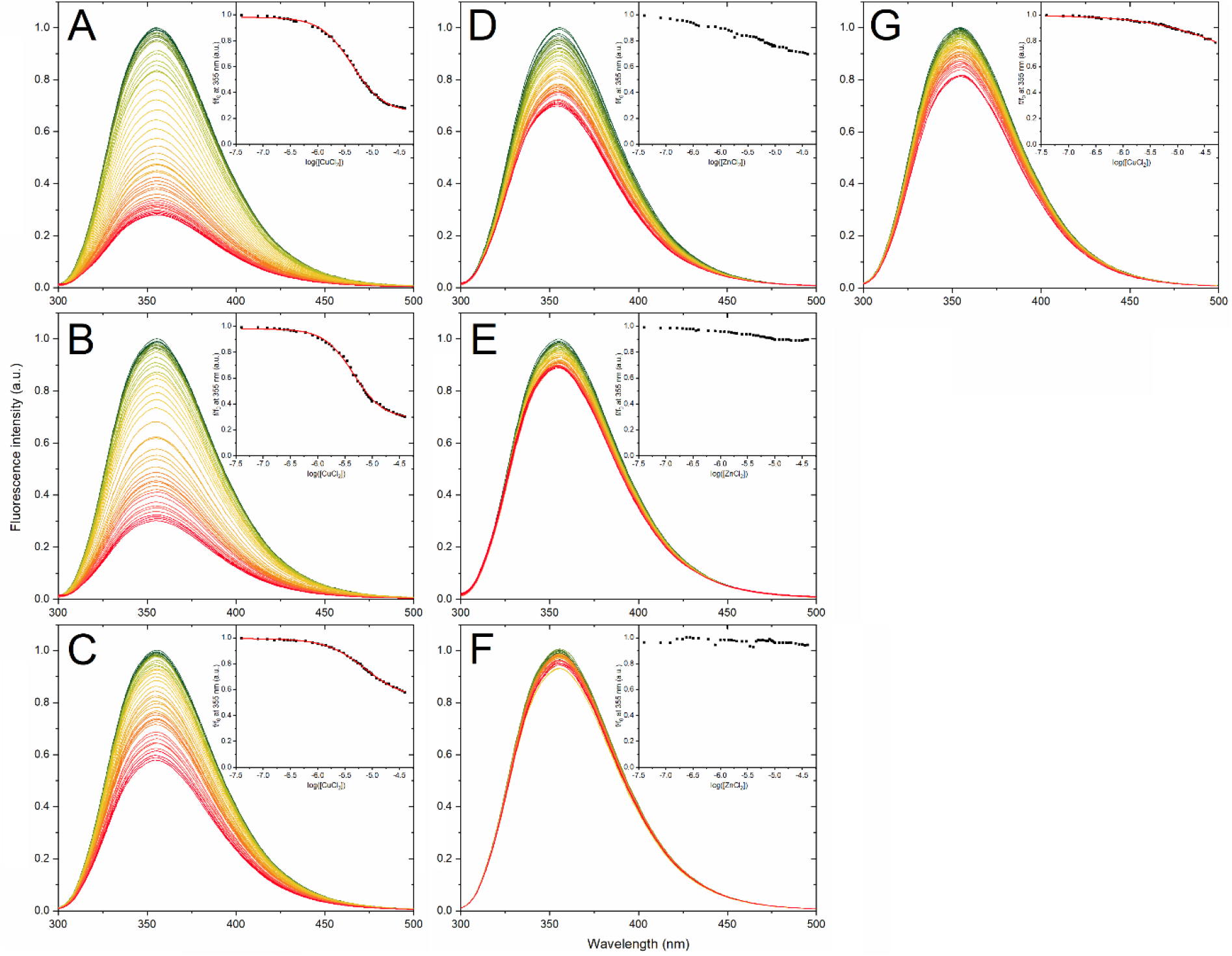
Fluorescence spectra for 5 μM OR peptide at 25 °C quenched with: (**A**) CuCl_2_ in 10 mM phosphate buffer, pH 7.5; (**B**) CuCl_2_ in 10 mM MES buffer, pH 7.5; (**C**) CuCl_2_ in 10 mM MES buffer, pH 5.5; (**D**) ZnCl_2_ in 10 mM phosphate buffer, pH 7.5; (**E**) ZnCl_2_ in 10 mM MES buffer, pH 7.5; (**F**) ZnCl_2_ in 10 mM MES buffer, pH 5.5; (**G**) CuCl_2_ in 1 mM TCEP, 10 mM MES buffer, pH 7.5.

To investigate if the OR peptide also binds Cu(I) ions, we performed tryptophan fluorescence titrations with CuCl_2_ under reducing conditions obtained with 1 mM TCEP. Our results showed that titrating Cu(I) ions to 5 μM OR peptide in 10 mM MES buffer, pH 7.5, and 1 mM TCEP, clearly quenched the tryptophan fluorescence (Fig. 5G), demonstrating binding also of Cu(I) to the OR peptide. Fitting the fluorescence data at 355 nm to eq. 2 yielded an apparent K_d_ of ~81 μM. As no inflection point was observed for the binding curve, this value should be considered a rough estimation. But even so, it appears that the OR peptide has weaker affinity for Cu(I) ions than for Cu(II) ions.

Titrations with Zn(II) ions to 5 μM OR peptide at 25 °C were performed in the same three buffers as the titrations with CuCl_2_, i.e. 10 mM phosphate buffer, pH 7.5; 10 mM MES buffer, pH 7.5 and 10 mM MES buffer, pH 5.5. The changes in the OR tryptophan fluorescence at pH 7.5 were stronger in the phosphate buffer than in the MES buffer, indicating a buffer effect on the zinc binding (Figs. 5D and 5E). At pH 5.5 no significant changes in fluorescence were observed (Fig. 5F), indicating that the OR peptide does not bind Zn(II) ions at acidic conditions. In all titrations the fluorescence maximum was at 355 nm and did not change its position. The weaker fluorescence quenching by Zn(II) ions compared to Cu(II) ions is to be expected, as Zn(II) ions are not paramagnetic. However, the observed quenching effect does show that also Zn(II) ions bind to the peptide at neutral pH, even though the signal-to-noise ratio is too low to quantitatively evaluate the binding curves.

### 3.4. Cu(II) and Zn(II) ions bound to the OR peptide both induce formation of hairpin structures

Molecular dynamics (MD) simulations were carried out to characterize the structural transitions in the OR peptide, i.e. PrP^C^(58-93), when bound to Cu(II) and Zn(II) ions. Final models from the MD simulations are shown in Figs. 6, 7, and 8, and visualizations of the first principal components of the simulations are shown in Fig. S3.

**Figure 6.**
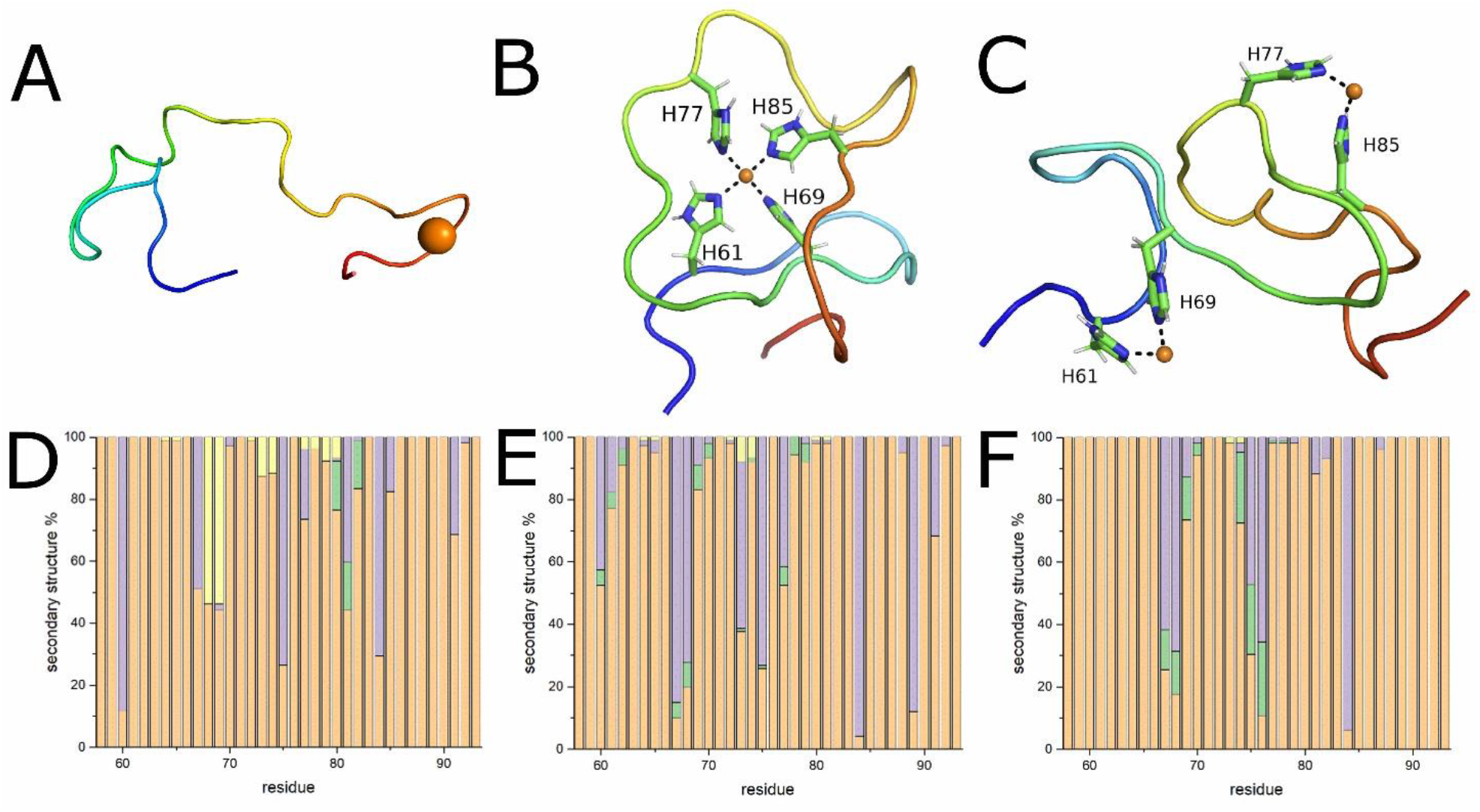
Final snapshots and secondary structure distributions of the OR peptide simulated with: (**A, D**): a single Cu(II) ion and protonated histidine N^ε2^ atoms; (**B, E**): a single Cu(II) ion and neutral histidine N^ε2^ atoms; (**C, F**): two Cu(II) ions and neutral histidine N^ε2^ atoms. The secondary structures were determined for each generated model using the PROSS^64^ algorithm: β-turns are shown in yellow, polyproline II helices in violet, β-strands in green, and coils in orange.

**Figure 7.**
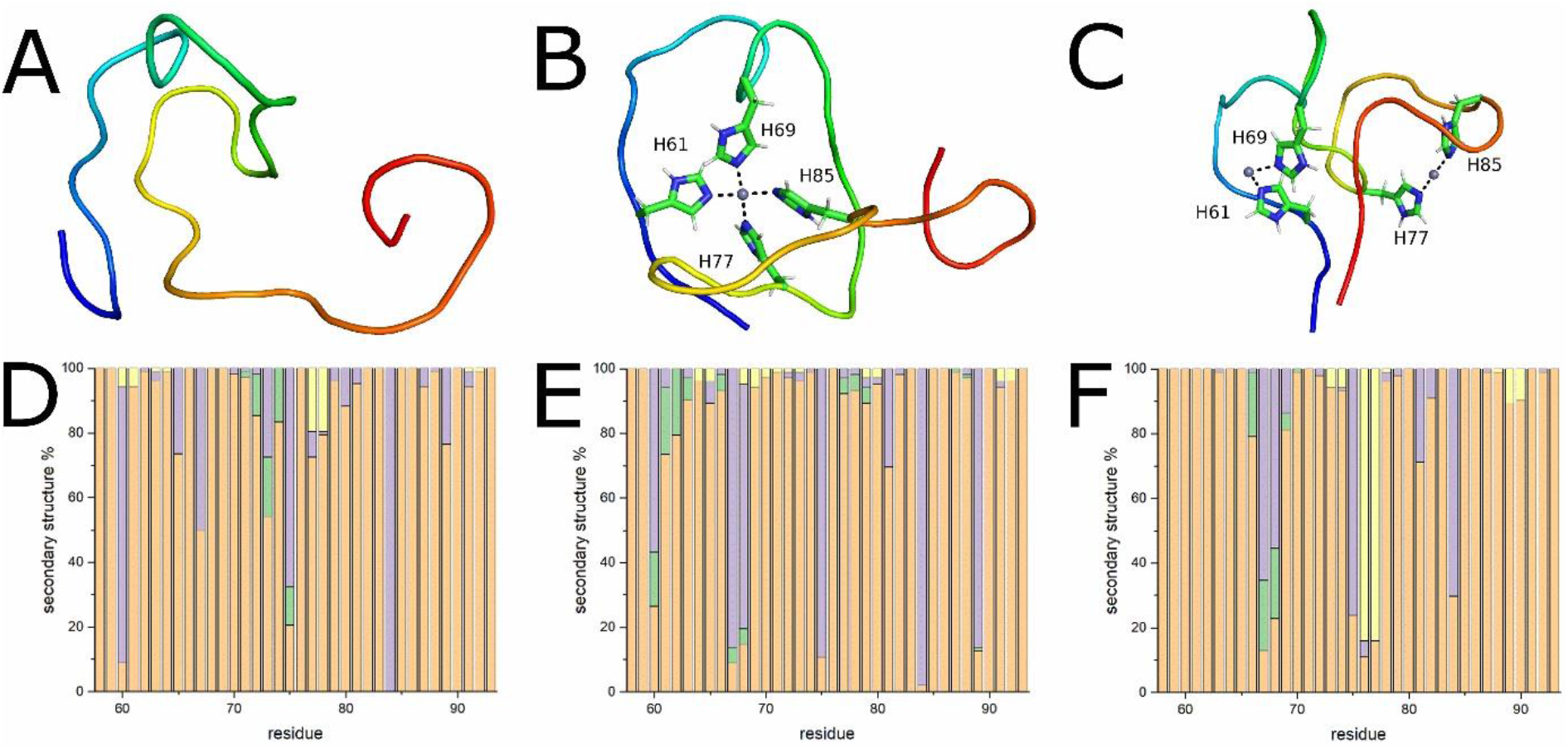
Final snapshots and secondary structure distributions of the OR peptide simulated with: (**A, D**): a single Zn(II) ion and protonated histidine N^ε2^ atoms; (**B, E**): a single Zn(II) ion and neutral histidine N^ε2^ atoms; (**C, F**): two Zn(II) ions and neutral histidine N^ε2^ atoms. The secondary structure distributions were calculated using the PROSS method: β-turns are shown in yellow, polyproline II helices in violet, β-strands in green, and coils in orange.

**Figure 8.**
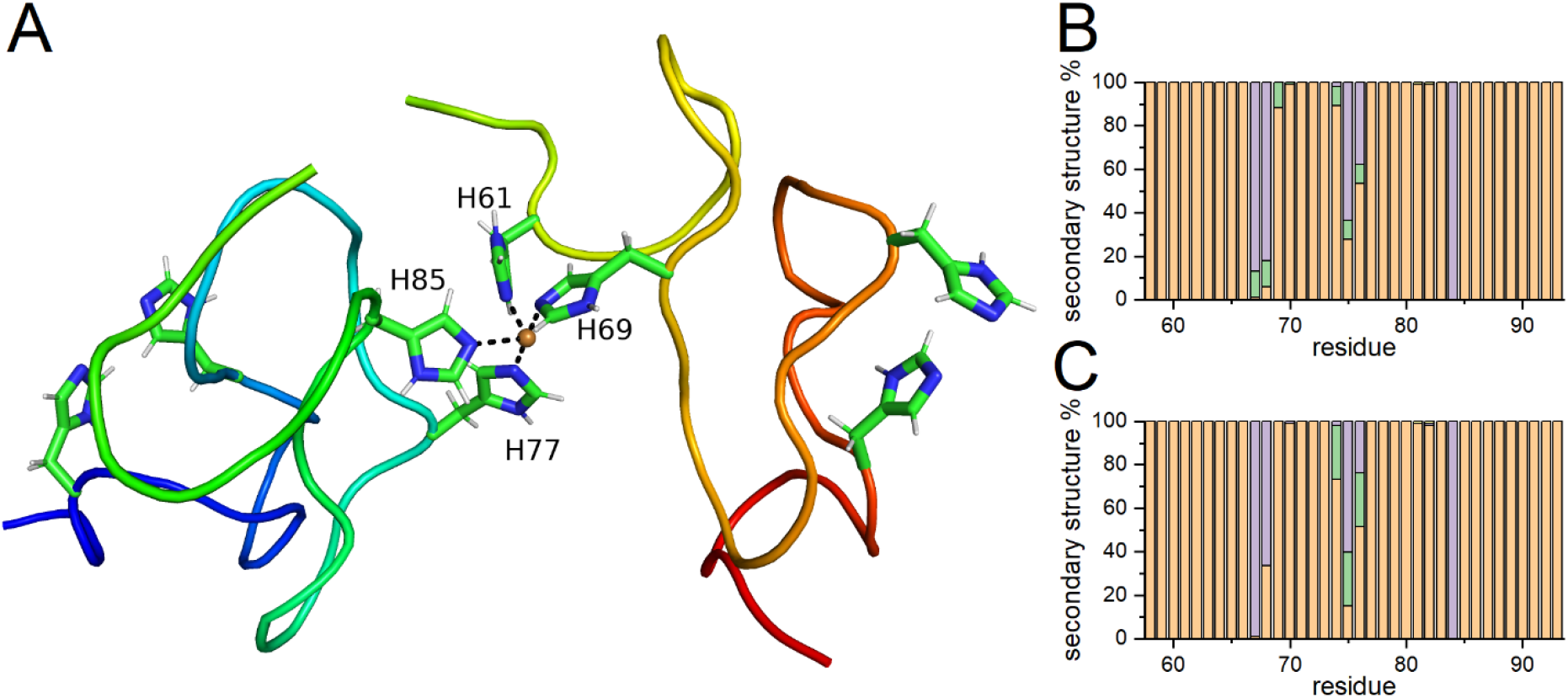
Final snapshot (**A**) of two OR peptide molecules simulated with a single Cu(II) ion. The N-terminus and C-terminus of the first OR peptide (OR-1) are marked in respectively blue and green, and the N-terminus and C-terminus of the second OR peptide (OR-2) are marked in respectively green and red. The secondary structure distributions for the two OR peptides are shown in (**B**) (OR-1) and (**C**) (OR-2), and were calculated using the PROSS method: β-turns are shown in yellow, polyproline II helices in violet, β-strands in green, and coils in orange.

To study interactions corresponding to acidic pH, a single Cu(II) ion was positioned next to four protonated histidine N^ε2^ atoms. As expected, the Cu(II) ion rapidly moved away from the protonated His residues during the MD equilibration phase. The simulations quickly converged, as shown by the RMSD values (Figure S4A). During the whole simulation time, the Cu(II) ion remained bound to the Cβ main chain carbonyl groups of residues His85, Gly87, Gly88, and Trp89, suggesting non-specific electrostatic binding (Fig. 6A). The OR peptide in this protonation state adopted an extended and flexible conformation with an RMSF around 0.5 nm (Figure S5A), where the secondary structure mainly was coil with some regions showing propensities for β-turn and PPII helix conformations (Fig. 6D). Principal component analysis (PCA) suggested that the main conformational changes involved motions in the N- and C-termini (Figure S3A).

Next, we simulated the OR peptide with either one or two Cu(II) ions located next to the neutral N^ε2^ atoms of the four OR histidine residues, corresponding to the OR peptide at neutral pH. Both simulations quickly converged (Fig. S4A), and the single Cu(II) ion remained bound by all four histidine residues and two axially-bound water molecules over the whole simulation time (Fig. S6A). In this binding mode the peptide backbone formed multiple loops around the Cu(II) ion, while residues His61-Gly71 formed a hairpin-like structure stabilized by Cu(II)-bound His61 and His69 (Fig. 6B). This model had smaller RMSF values than the OR peptide model with protonated histidine residues (Fig. S5A), indicating a more rigid structure. Indeed, the main PCA component showed smaller backbone displacements near the Cu(II) ion than at the N- and C-termini (Figure S3B). Moreover, in this binding mode more OR residues adopted PPII helix and transient β-strand secondary structures, compared to the OR peptide with protonated histidine residues (Fig. 6E). For the simulations with two Cu(II) ions, each copper ion remained bound by two histidine residues and four water molecules (Fig. 6C) over the whole simulation time (Fig. S6B). In this binding mode, the peptide backbone formed a structure with three hairpins. The first hairpin was stabilized by one of the Cu(II) ions, and was located near the N-terminus, involving residues Gly62-His69. The second hairpin, which was stabilized by the other Cu(II) ion, involved residues Gly74-His85 and was thus located near the C-terminus. The third hairpin involved residues His69-Pro76 and formed a bridge between the two Cu(II)-bound segments (Fig. 6C). The RMSF data showed intermediate values, with minima corresponding to the histidine residues involved in Cu(II)-binding (Fig. S5A). The primary PCA component suggests that the two Cu(II)-stabilized hairpin structures were rigid in themselves, but could move relative to each other, resulting in formation of the middle bridging hairpin (Fig. S3C). Secondary structure analysis indicated a reduction of PPII structure and formation of antiparallel β-strands around residues Gln67-Gly70 and Gly74-Pro76, i.e. roughly the region of the middle bridging hairpin (Fig. 6F).

Similar simulations were performed also for the OR peptide with Zn(II) ions. For the OR peptide with fully protonated histidine residues and simulated with a single Zn(II) ion, the metal ion moved away from the peptide in the MD equilibration phase and remained unbound for the whole simulation time. The peptide adopted an elongated (Fig. 7A) and flexible conformation with an average RMSF of 0.6 nm (Fig. S5B). The first PCA component suggests that the main movement in the OR peptide involves motions in the N- and C-termini (Fig. S3D). During the simulation time the OR peptide here adopted mainly coil and PPII secondary structures (Fig. 7D).

In the next step we simulated the OR peptide with neutral N^ε2^ atoms, together with one or two Zn(II) ions. Both simulations converged (Fig. S4B). The single Zn(II) ion placed next to the four histidine residues remained bound over the whole simulation time (Fig. S6C) and was additionally coordinated by two axially bound water molecules. In this binding mode the peptide backbone again formed multiple loops around the Zn(II) ion, and residues Gly74-Gly86 formed a hairpin structure stabilized by the zinc ion bound to His77 and His85. The average value of the Cα RMSF was 0.3 nm, which is a much smaller value than for the OR peptide with protonated histidine residues (Fig. S4B). The first PCA component suggests that the main movement of the peptide backbone occurred in the hairpin loop (residues Gly78-Gly82) and the C-terminal loop (residues 86-91), while the backbone around the Zn(II) ion had smaller mobility (Fig. S3E). Analysis of the secondary structure showed that more residues formed a β-strand secondary structure than for the OR peptide with protonated histidine residues (Fig. 7E). However, the peptide mainly adopted coil and PPII helix secondary structures.

When two Zn(II) ions were positioned close to the histidine residues, both ions again remained bound over the whole simulation time, and each was coordinated by two histidine residues and four water molecules (Fig. S6D). In this binding mode the peptide backbone formed four hairpin structures. Two hairpin structures, located at the N- and C-termini, involved respectively residues His61-His69 and His77-His85, and each of them was stabilized by a single Zn(II) ion. Then, two bridging hairpins involving respectively residues Gln67-Trp73 and Gln75-Gly78 were located between the two Zn(II)-stabilized hairpins (Fig. 7C). The peptide backbone had an average RMSF of 0.4 nm (Fig. S5B), indicating higher Cα fluctuations than when only one Zn(II) ion was bound to the OR peptide. The primary PCA component suggests that in this binding mode, the main motion of peptide backbone corresponds to formation of the two bridging hairpin structures (residues Gln67-Trp73 and Gln75-Gly78) and closing the whole model by terminal residues Gly58, Gln59 and Gly92, Gly93 (Fig. S3F). Analysis of the peptide’s secondary structure showed a reduced PPII helix content, but an increase in the coil and β-turn secondary structures (Fig. 7F).

In the last step we simulated two OR peptide molecules with neutral N^ε2^ atoms, bound to a single Cu(II) ion. The initial model contained the Cu(II) ion bound to His77 and His 85 from the first OR molecule (OR-1), and to His61 and His69 from the second OR molecule (OR-2). The models of OR-1 and OR-2 were taken from the last MD step of our OR simulation together with two Cu(II) ions. The simulation converged and surprisingly showed a minimal RMSD over time (Fig. S4A). The average value of the Cα RMSF was below 0.05 nm (Fig. S5C), which is the smallest value from all simulations performed in this study, and the single Cu(II) ion placed next to the four histidine residues remained bound during the entire simulation time (Fig. S6E). The three hairpin structures, previously formed in the OR peptide simulated with two Cu(II) ions, were preserved in each OR molecule (Fig. 8A). Secondary structure analysis indicated similar structural compositions for the OR-1 and OR-2 molecules. The two peptide molecules predominantly adapted coil structures, with a PPII structure at residues Gln67, Pro68, Gln75, Pro76, Pro84 in OR-1 and OR-2, and β-strands at residues Gln67-His69, Gly74-Pro76 in OR-1, and Gly74-Pro76 in OR-2 (Fig. 8B, C).

### 3.5 Calculation of pKa values for the OR peptide histidines

Protein binding affinities for Cu(II) and Zn(II) ions can be affected by the pKa values of the His residue side chains, which are known to be close to the physiological pH^59,82^. Changes in pKa values can be observed in structural conformations that favor hydrogen bond interactions with the histidine side chains, and such pKa changes can affect the binding affinity for metal ions.

We therefore calculated pKa values for His residue side chains for different MD conformations (Table 2). To increase the accuracy of the calculations, two independent protocols were used, namely PropKa 2.0 and DelPhiPKa^66,68^. The two protocols gave pKa values for all four His residues that are maximally within ± 6% of the average calculated values (Table 2).

**Table 2.**
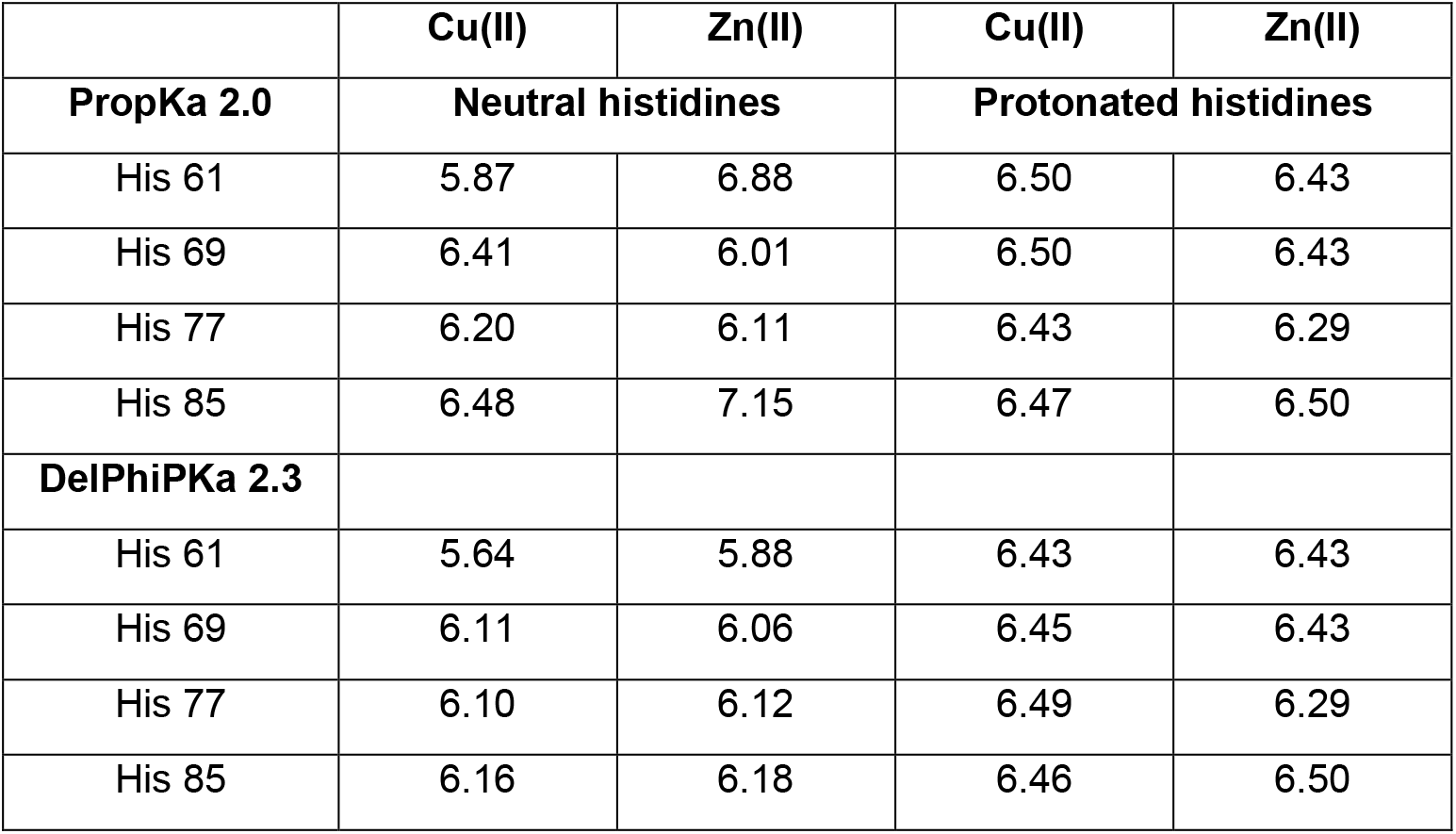
pKa values for the four histidine residues in the OR peptide, calculated for both neutral and protonated histidine, and with either the PropKa 2.0 or the DelPhiPKa 2.3 protocol.

|  | Cu(II) | Zn(II) | Cu(II) | Zn(II) |
| --- | --- | --- | --- | --- |
| **PropKa 2.0** | **Neutral histidines** |  | **Protonated histidines** |  |
| His 61 | 5.87 | 6.88 | 6.50 | 6.43 |
| His 69 | 6.41 | 6.01 | 6.50 | 6.43 |
| His 77 | 6.20 | 6.11 | 6.43 | 6.29 |
| His 85 | 6.48 | 7.15 | 6.47 | 6.50 |
| **DelPhiPKa 2.3** |  |  |  |  |
| His 61 | 5.64 | 5.88 | 6.43 | 6.43 |
| His 69 | 6.11 | 6.06 | 6.45 | 6.43 |
| His 77 | 6.10 | 6.12 | 6.49 | 6.29 |
| His 85 | 6.16 | 6.18 | 6.46 | 6.50 |

The two protocols consistently show that both the position of the His residue and the protein conformation have noticeable effects on the pKa values (Table 2). For the final models from the MD simulations, both protocols produce slightly higher pKa values for His 85 than for the other three residues. His 85 is the histidine most exposed to the solvent, and ionic interactions with water molecules are energetically more favorable than hydrogen bonds. For the same reasons, both protocols consistently show slightly higher pKa values for the final models from simulations with charged His residues, than for models from simulations with neutral His residues.

## 4. Discussion

### 4.1 Solution structure of the OR peptide

Previous studies have suggested that the OR peptide in aqueous solution adopts a combination of random coil structure together with either PPII left-handed extended helix^70–72^ or β-turn structures^35,73,74^. Our CD results for the apo-OR peptide, i.e. PrP^C^(58-93), show unfolding at elevated temperatures (Fig. 2), thus demonstrating the existence of secondary structures different from random coil at low temperatures. In the 20-65 °C temperature range an isodichroic point at 204 nm indicates a PPII-helix to random coil transition^76^. This is consistent with previous studies of the OR peptide ^70–72^, and also similar to our earlier results on the structure of the Aβ peptides involved in Alzheimer’s disease^75^. Temperature studies of the Aβ(1-40) peptide and its shorter N-terminal fragments generally show an isodichroic point around 208 nm, a weak positive band at ~222 nm that becomes negative at high temperatures, and a strong negative band at ~200 nm whose intensity is reduced at high temperatures. The similar results obtained here for the OR peptide suggest similar structures and temperature-induced structural transitions in the two peptides. Our results suggest that ~45% of the OR peptide is in PPII conformation at 37°C, with the remaining structure being random coils. For the Aβ(1-40) peptide, the corresponding numbers are ~30% PPII helix and ~70% random coil structure^75^. The lack of a well-defined isodichroic point below 20 °C (Fig. 2) indicates that the OR peptide can form various secondary structures at low temperatures. This is consistent with NMR studies of the OR sequence at 20 °C, which suggest the presence of structured loops and β-turns^83^.

### 4.2 Metal binding to the OR peptide induce β-sheet formation

Earlier studies have shown that the OR domain can bind up to four Cu(II) ions, but no detailed structural model for the OR peptide backbone during Cu(II) binding has been proposed. More than twenty years ago Viles et al. performed far-UV CD titrations with Cu(II) ions to the OR peptide in water at pH 7.4^35^. Addition of Cu(II) ions was found to decrease the intensity of both a negative band at 200 nm and a positive band at 225 nm, and to induce a new negative band at 222 nm together with a new positive band at 204 nm. This was interpreted as a structural alteration corresponding to formation of β-turns or structured loops^35^. Later studies have suggested that the negative band at 222 nm might reflect the structures of tryptophan side chains^27^. In this study we were able to largely recreate the results of Viles et al., but we present a different interpretation. The difference spectrum for the Cu(II)-OR complex in water (Fig. S2A) has a strong negative band at 222 nm and a strong positive band at 202 nm. It thereby resembles the CD spectrum for left-handed twisted antiparallel β-sheets^77^, as well as the CD spectrum for a hydrophobic fragment of the Aβ peptide, i.e. Aβ(25-35), to which we previously have attributed an antiparallel β-sheet secondary structure^75^. We therefore argue that binding of Cu(II) ions to the OR peptide in water results in formation of antiparallel β-sheet structures.

We further propose that antiparallel β-sheets are formed also when Cu(II) ions bind to 5 μM OR peptide in the phosphate buffer, pH 7.5, but these β-sheets then appear to have a different geometry. Addition of up to 5 μM CuCl_2_, corresponding to a 1:1 OR:Cu(II) molar ratio, produces an isodichroic point at 217 nm, which suggests a two-state PPII-helix to β-turn transition. Above 5 μM of CuCl_2_, the new CD band at 208 might be caused by formation of a relaxed antiparallel β-sheet secondary structure^77^. The intensity increase for the new 208 nm band was higher for Cu(II) concentrations in the 5 μM to 10 μM than for the 10 μM to 40 μM interval, suggesting that binding of two Cu(II) ions to the OR peptide is a primary event. Further binding up to four Cu(II) ions increases the formation of antiparallel β-sheet structure, however to a lesser amount.

In an earlier study, a CD spectrum for 50 μM of Syrian hamster OR peptide together with 250 μM of Cu(II) ions in 20 mM ammonium acetate at pH 6.0^84^, is almost identical to our CD spectrum for 5 μM OR peptide with 10 μM of Cu(II) ions in 10 mM phosphate buffer, pH 7.5 (Fig. 3B). The CD spectrum of the Syrian hamster OR peptide in the ammonium acetate buffer was interpreted as representing a Cu(II)-OR complex^84^. The CD spectrum of this proposed Cu(II)-OR complex was different than for the previously reported complex in water at pH 7.4^35^, which is consistent with our current observations (Figs. 3A and 3B). In another study, no changes in the CD spectrum were reported when Cu(II) ions were titrated to the OR peptide in 10 mM phosphate buffer, pH 7.0^72^. Although the OR peptide concentration was not specified and CD data were shown only for one CuCl_2_ concentration, the reported CD spectra look like they might have an isodichroic point around ~220 nm, which is close to our observed isodichroic point at 217 nm (Fig. 3B). The authors explained the lack of a clear effect when adding copper ions by phosphate ions being able to compete for Cu(II) binding^72^. In living organisms Cu(II) ions never exist in free form^32^, and phosphate ions may indeed form complexes with Cu(II) ions^85^. If binding of Cu(II) ions to phosphate was so much stronger than binding to the OR peptide, then Cu(II) binding to the OR peptide would be physiologically irrelevant, given the high concentration of free phosphate ions in the extracellular fluid^86^. But our current data clearly show that the OR peptide binds both Cu(II) and Zn(II) ions, also in the phosphate buffer.

As the OR domain is known to bind multiple - up to four - Cu(II) ions, the binding modes depend on the Cu(II) concentration^37,73,78,80^. The first Cu(II) ion is coordinated by four N^ε2^ atoms from the four histidine residues^37,80^. This binding mode has the highest affinity, with a K_d_ below 3 nM^38,80^. With two Cu(II) ions, each one is coordinated by two histidine sidechain N^ε2^ atoms^37^, possibly together with negatively charged atoms, and the K_d_ is ~200 nM^80^. Further addition up to four Cu(II) ions rearranges the binding configuration, so that each Cu(II) ion is coordinated by one histidine sidechain N^δ1^ atom and three negatively charged atoms, such as two deprotonated amide nitrogens and one carbonyl oxygen from the preceding glycine residues^36,37,80^. In this binding mode the OR peptide has the weakest affinity for Cu(II) ions, with a K_d_ in the 1 – 10 μM range^38,80^.

The OR peptide backbone is expected to adopt different conformations for each binding configuration. Our CD titrations confirm this notion (Fig. 3B), and the transitions are particularly clear at 224 nm (Fig. 4B). Up to 5 μM CuCl_2_, i.e. 1:1 Cu(II):OR ratio, the intensity at 224 nm monotonically decreased. From 5 μM to 10 μM CuCl_2_, i.e. 1:1 to 2:1 Cu(II):OR ratio, the intensity at 224 nm monotonically increased. From 10 μM to 40 μM CuCl_2_, i.e. 2:1 to 8:1 Cu(II):OR ratio, the intensity at 224 nm decreased and reached a plateau around ~20 μM CuCl_2_, i.e. around 4:1 Cu(II):OR ratio. These observations are clearly consistent with binding of one, two, and up to four Cu(II) ions to the OR peptide in the three intervals of CuCl_2_ concentration (Fig. 4B).

Binding of Zn(II) ions to the OR peptide in 10 mM phosphate buffer, pH 7.5, seems to induce similar structures in the OR peptide as Cu(II) ions. However, while binding of Zn(II) ions also appears to induce formation of an antiparallel β-sheet secondary structure, the geometry of this β-sheet could be different^77^. Again, the changes in CD intensity at 224 nm seem to reflect distinct structural transitions (Fig. 3C). For additions of up to 5 μM of ZnCl_2_, i.e. 1:1 Zn(II):OR ratio, the 224 nm intensity monotonically decreased, suggesting binding of a single Zn(II) in this interval (Fig. 4D). From 5 μM to 10 μM of ZnCl_2_, i.e. from 1:1 to 2:1 Zn(II):OR ratios, the 224 nm intensity decreased even more steeply, suggesting binding of two Zn(II) ions in this interval. Above 10 μM ZnCl_2_ no further changes appear in the CD spectrum at 224 nm, indicating that binding of two Zn(II) ions saturates the OR peptide. Other researchers have suggested that all four OR histidine residues are involved in binding a single Zn(II) ion^47,48,87^. This might be true when the OR peptide binds Zn(II) ions in a 1:1 molar ratio, but at higher Zn(II) concentration we suggest that two Zn(II) ions are bound by two histidine residues each, in a similar fashion as for Cu(II) ions (above).

### 4.3 Histidine protonation and metal ion binding to OR peptide

Our fluorescence measurements of the OR peptide titrated with Cu(II) ions at pH 7.5 suggest an apparent K_d_ ~4.5 μM (Table 1). This value is close to the mean apparent dissociation constant of 7.2 μM reported for an isolated HGGGW repeat^38^. Calculating real K_d_ values requires competition experiments with e.g. glycine^38^. As our main objective was to show changes in metal ion binding affinity under different conditions, no competition experiments were performed in this study. Our CD experiments for the OR peptide titrated with Cu(II) ions clearly shows three spectral transitions, that we attribute to binding of four Cu(II) ions. The single apparent K_d_ value from our fluorescence experiments therefore seems to represent the averaged K_d_ from all three possible Cu(II) binding modes^80^. In our fluorescence experiments under acidic conditions at pH 5.5 (Fig. 5C), the K_d_ value (~7.4 μM) suggested slightly weaker binding than at pH 7.5 (~4.5 μM) (Table 1). Despite this small difference in binding affinity, this result clearly shows that the OR peptide can bind Cu(II) ions also under acidic conditions where the histidines are protonated^59^ as their pKa values are between 6 and 7, depending on the degree of solvent exposure and hydrogen bonds created (Table 2). Earlier studies with mass spectrometry indicate that the OR peptide can bind only up to two Cu(II) ions at pH 6.0, i.e. less than the four Cu(II) ions that can be bound at pH 7.4^84^. Thus, our fluorescence quenching measurements at pH 5.5 (Fig. 5C) may reflect the average affinity for binding two Cu(II) ions to OR peptide. Tryptophan fluorescence measurements of the OR peptide with ZnCl_2_ at pH 7.5 showed reduced fluorescence intensity indicative of binding, but the poor signal-to-noise ratio of the data did not allow the binding affinity to be calculated (Figs 5D and 5E). At pH 5.5 addition of ZnCl_2_ did not affect the intrinsic OR peptide fluorescence at all (Fig. 5F), indicating that the OR peptide does not bind Zn(II) ions under acidic conditions.

### 4.4 Models for β-sheet formation

Molecular dynamics simulations of the OR peptide with metal ions generally support our CD results. The secondary structure of the OR peptide with protonated histidine residues, in the presence of a single Cu(II) or Zn(II) ion, mainly consists of coil structure, although some PPII-helix structure (12% overall) is present around residues Pro60, Gln67, Gln75 and Pro84 (Fig. 6D, Fig. 7D). When the OR peptide with neutral histidine residues is modeled together with a single bound metal ion, the peptide again mainly formed a coil structure, although together with four transient β-strands. For a bound Cu(II) ion, the transient β-strands were Pro60-Gly62, Gln67-Gly70, Trp73-Gln75, and His77-Gly79. For a bound Zn(II) ion, the transient β-strands were Pro60-Gly63, Gly66-Pro68, His77-Gln79, and Gly87-Trp89. Thus, most of the β-strands appeared in close proximity to the histidine residues involved in metal ion binding, i.e. His61, His69, His77, and His85. The overall amount of PPII-helix structure was around 15% in these simulations, with the main contributions coming from Pro60, Gln67, Pro68, Gln75, Pro84 and Trp89.

Although the OR peptide simulated with neutral histidine residues and two bound metal ions mainly adopted a coil structure, we observed significant reduction in the PPII-helix structure (9% overall), which is in line with our CD analysis. The PPII-helix content was here distributed in a more uniform way, mainly around the Pro68, Pro76, and Pro84 residues. Furthermore, modeling with two bound Cu(II) ions (Fig. 6C) induced formation of two β-strands consisting of residues Gln67-Gly70 and Gly74-Pro76, i.e. mainly in the middle of the bridging hairpin structure (residues His69-Pro76). For two bound Zn(II) ions, a similar β-strand was formed at Gly66-Gly70, together with two accompanying short β-turns at Trp73-Gly74 and Pro76-His77. In this case the formed β-structures were located around the first (residues Gln67-Trp73) and second (residues Gln75-Gly78) bridging hairpins, which connect the N-terminal and C-terminal hairpin structures stabilized by the two metal ions (Fig. 7C). These observations are consistent with our CD results, where formation of antiparallel β-sheets was observed for the second spectral transition (i.e., for binding of the second Zn(II) ion (Fig. 4D).

Structural models with a single bound metal ion from our MD simulations agree with previously published results for a single Cu(II) ion bound to the OR peptide, where the metal binding site remained exposed to the solvent and the peptide backbone formed multiple loops around the Cu(II) ion^88,89^. Interestingly, Pushie et al. performed molecular dynamics simulations for two octapeptide repeats, each bound with a single Cu(II) ion^90^. The authors observed stacking of the two HGG metal binding regions on each other, where the GWGQ linker region formed a bend or a turn structure. Those models resemble our hairpin structures from the simulation of the OR peptide with two Cu(II) ions. In the “stacked” models two copper binding sites were located in close proximity, with a copper-copper distance of ~5 Å. With two neighboring histidine imidazole moieties, it is possible that the 2:1 binding mode (two Cu(II) ions/one OR peptide) under higher Cu(II) occupancy undergoes rearrangement to a 4:1 binding mode, with the hairpin properties being preserved, which would agree with our CD results. Future modeling with multiple OR peptides (or full-length PrP^C^) and multiple metal ions may shed more light on the structural effects of metal binding to aggregated peptides/proteins.

### 4.5 OR peptide metal ion binding and amyloid formation

The OR domain is an important part of PrP^C^, being involved in the protein’s neuroprotective effect^91^. Previous studies have suggested that Cu(II) and Zn(II) ions may inhibit in vitro conversion of PrP^C^ to PrP^Sc^ by formation of nonamyloid aggregates^92^. However, we have previously reported that after addition of four molar equivalents of Zn(II) ions, the OR peptide forms structured fibrils that bind the amyloid-specific dyes Thioflavin T and Congo Red, and which display the cross-β structure typical for amyloid material ^50^. Thus, the OR peptide incubated with an excess of Cu(II) ions may also form amyloid aggregates. Our CD titrations and MD simulations of the OR peptide bound with two metal ions (i.e., Cu(II) or Zn(II)) showed formation of β-structures, which is consistent with our previous results. In our MD simulations the OR peptide formed β-structures in the Gln67-Gly70 and Gly74-Pro76 regions, and as β-structures – especially hairpins – are favorable for amyloid formation^6^, we speculate that these regions may form a core for amyloid aggregation of the OR peptide or full-length PrP^Sc^. However, our CD and MD results indicate that binding of a single metal ion to the OR peptide induces only minor changes in the peptide’s secondary structure. Thus, it is possible that only high concentrations of metal ions – more than 1:1 ratio – would induce secondary structures suitable for amyloid formation. On the other hand, binding of divalent metal ions to full-length PrP^C^ is known to induce interactions between the metal-bound OR domain and the C-terminal helical domain^25,87,93^. Such interactions likely help to hold the protein structure together, and it is generally known that structured proteins first must become unstructured before they can “misfold” into β-sheets (or β-sheet hairpins) and begin to assemble into amyloid forms. In a PrP^C^ variant with point mutations, corresponding to genetic Creutzfeldt-Jakob disease, fatal familial insomnia, and the Gerstmann-Straussler-Scheinker disease, addition of Zn(II) ions induced broadening of NMR peaks indicative of weaker interactions between the Zn(II)-bound OR region and the C-terminal domain^87^ than in the native protein. Thus, the metal-induced interactions between the OR region and the C-terminal domain likely counteracts amyloid formation, by stabilizing the protein fold, and these stabilizing interactions may be weaker in some disease-related PrP^C^ mutants. Finally, metal ions may promote aggregation by binding to the OR region of two or more PrP^C^ proteins, thereby bringing the two proteins together (a first step towards aggregation). Our MD simulations (Fig. 8) showed that such intermolecular Cu(II) coordination was very stable over time, indicating that complexes with Cu(II) ions and two or more PrP^C^ molecules are likely to form *in vivo*.

The relation between metal binding and PrP^C^ aggregation is clearly complex, with two effects that likely promote aggregation (one metal ion binding multiple PrP^C^ molecules, and metal ions inducing β-sheet structures suitable for amyloid formation), and one effect that likely counteracts amyloid formation (stabilizing the protein fold by interactions between the C-terminal domain and the metal-bound OR region). Our tentative understanding is that the effects that promote aggregation will dominate at high concentrations of metal ions, i.e. at metal:protein ratios higher than 1:1. As four Cu(II) ions but only two Zn(II) ions could bind to one OR peptide, and as Cu(II) ions had a larger effect on the peptide structure than the Zn(II) ions, amyloid formation is probably more efficiently induced by Cu(II) than Zn(II) ions.

## 5. Conclusions

In summary, our results show that the OR region in the PrP^C^ protein can bind up to four Cu(II) ions or two Zn(II) ions. The average apparent binding affinities are in the low micromolar range for both metal ions (Table 1). The OR histidine residues are important binding ligands, where Zn(II) binding is more sensitive to histidine protonation than Cu(II) binding, and the metal ions can be coordinated by histidines from different PrP^C^ molecules – such intermolecular complexes appear to be stable first steps towards protein aggregation. Without bound metal ions, the secondary structure of the OR peptide is a combination of random coil and PPII helix. Addition of metal ions induces structural changes into β-sheet conformations, which generally are beneficial for amyloid aggregation. The structural conversions are most prominent for large concentrations (i.e. above 1:1 ratio) of Cu(II) ions, suggesting that especially Cu(II) ions could be an important factor in converting the PrP^C^ protein into amyloids of the neurotoxic PrP^Sc^ form.

## Acknowledgments

This study was supported by a research grant (2014/15/B/ST4/04839) from the National Science Centre in Poland to MK, by a grant from the Magnus Bergvall foundation in Sweden to SKTSW, and from grants from the Swedish Research Council and the Brain Foundation in Sweden to AG. High-performance computing at the University of Rijeka was supported by grants to ŽS from the European Fund for Regional Development (ERDF) and from the Ministry of Science, Education and Sports of the Republic of Croatia under project number RC.2.2.06-0001.

## Supporting material

**Figure S1.**
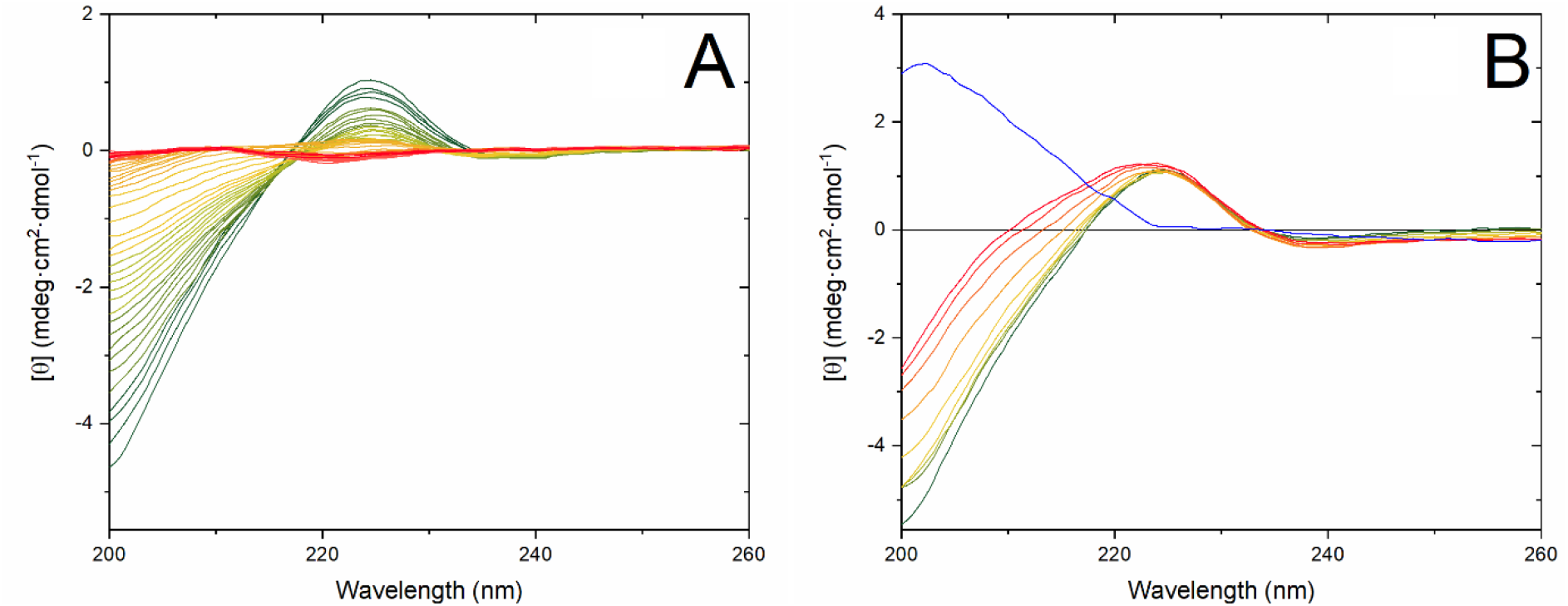
CD spectra of the OR peptide titrated with CuCl_2_. (**a**) Titration of 5 μM OR peptide in pure water at 25 °C, with CuCl_2_ in increasing (0.04 μM, 0.2 μM, 0.5 μM, 2 μM and 4 μM) intervals. The pH was adjusted to 7.5 with small amounts of NaOH. Initial spectrum: blue. Final spectrum (40 μM CuCl_2_): red. (**b**) Titration of 5 μM OR peptide in 10 mM phosphate buffer, pH 7.5, with CuCl_2_ in 5 μM intervals. Initial spectrum: green. Final spectrum (40 μM CuCl_2_): red. Difference spectrum between the initial and final states: blue.

**Figure S2.**
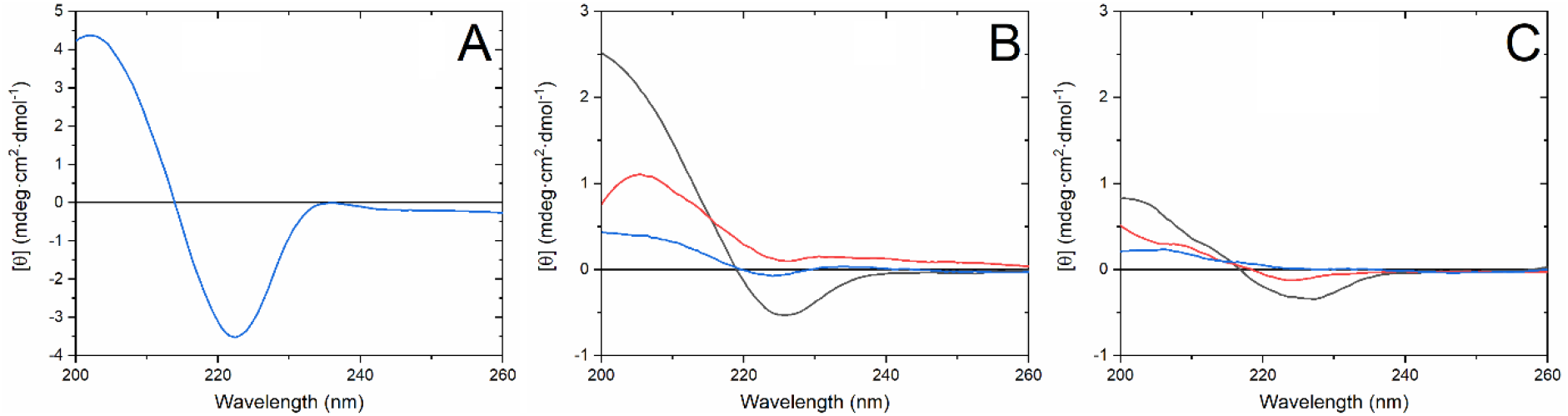
Difference spectra for CD curves in Figure 3. (**a**) Difference between the two spectra shown in Fig. 3a, i.e., 5 μM OR peptide in water at pH 7.5 before and addition of 20 μM CuCl_2_. (**b**) Difference spectra between the first and last spectrum of the first (black - 5 μM OR peptide with 5 μM CuCl_2_ minus 5 μM OR peptide), second (red - 5 μM OR peptide with 10 μM CuCl_2_ minus 5 μM OR peptide with 5 μM CuCl_2_), and third (blue - 5 μM OR peptide with 20 μM CuCl_2_ minus 5 μM OR peptide with 10 μM CuCl_2_) transitions shown in Figure 3b; (**c**) Difference spectra between the first and last spectrum of the first (black - 5 μM OR peptide with 5 μM ZnCl_2_ minus 5 μM OR peptide), second (red - 5 μM OR peptide with 10 μM ZnCl_2_ minus 5 μM OR peptide with 5 μM ZnCl_2_), and third (blue - 5 μM OR peptide with 20 μM ZnCl_2_ minus 5 μM OR peptide with 10 μM ZnCl_2_) transitions shown in Figure 3c.

**Figure S3.**
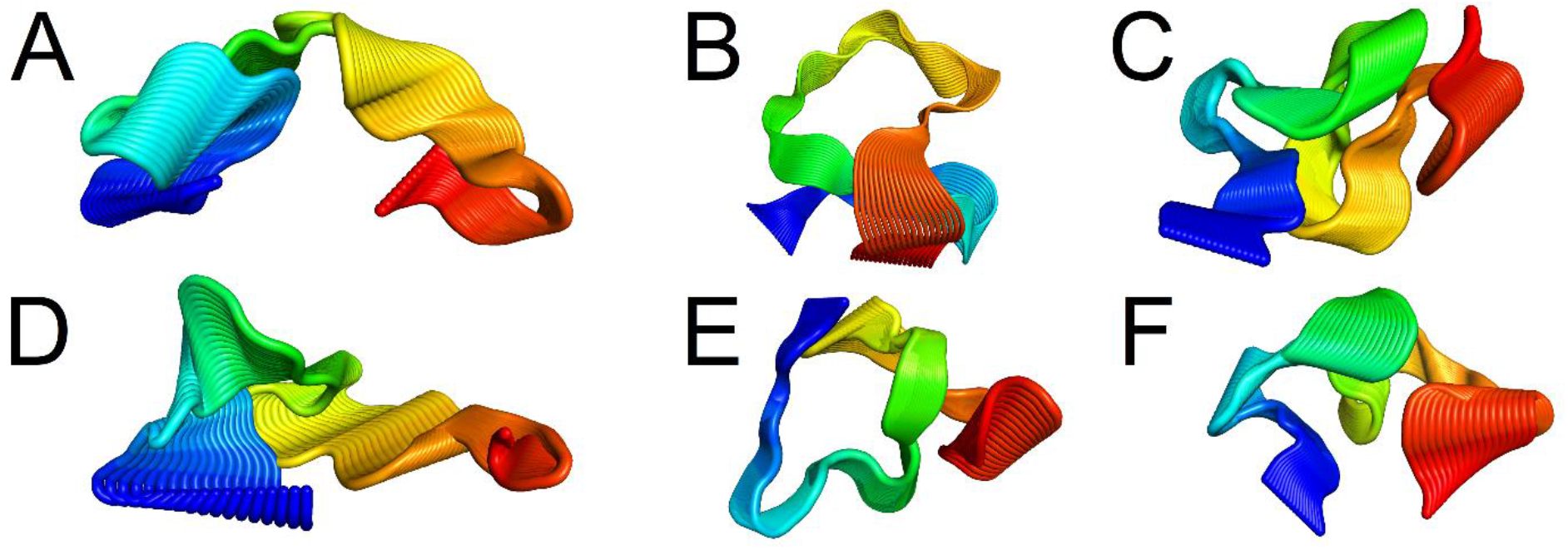
Visualization of the first principal component of the OR peptide simulated together with (**a**) a single Cu(II) ion and protonated histidine residues, (**b**) a single Cu(II) ion bound to four N^ε2^ atoms of neutral histidine residues, (**c**) two Cu(II) ions, each bound to two N^ε2^ atoms of neutral histidine residues, (**d**) a single Zn(II) ion and protonated histidine residues, (**e**) a single Zn(II) ion bound to four N^ε2^ atoms of neutral histidine residues and (**f**) two Zn(II) ions, each bound to two N^ε2^ atoms of neutral histidine residues.

**Figure S4.**
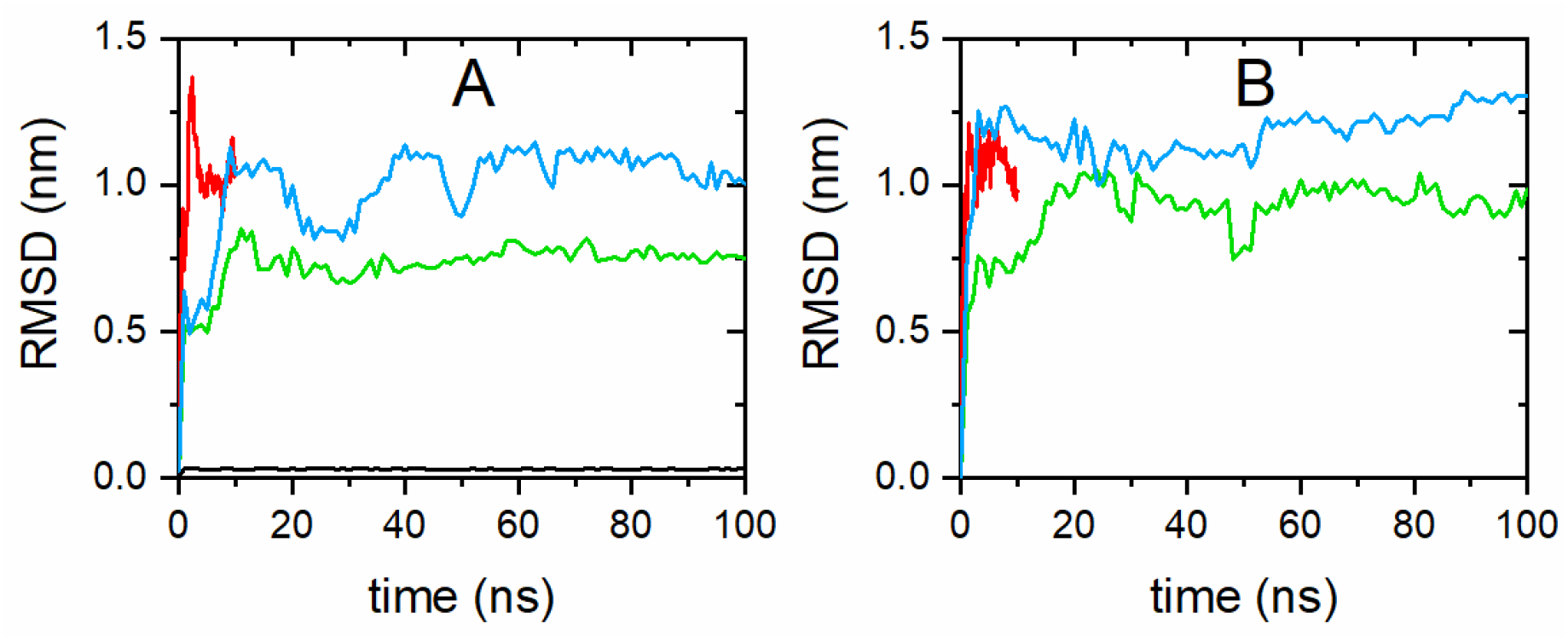
Root mean square deviation (RMSD) of atomic positions for simulations of the OR peptide together with: (**a**) Cu(II) ions, and (**b**) Zn(II) ions. The OR peptide with fully protonated histidine residues and a single metal ion (i.e., either Cu(II) or Zn(II)) is shown in red, while the OR peptide with neutral histidine residues simulated with a single metal ion is shown in green, the OR peptide with neutral histidine residues simulated with two metal ions is shown in blue, and the two OR peptides with neutral histidine residues simulated with a single metal ion are shown in black.

**Figure S5.**
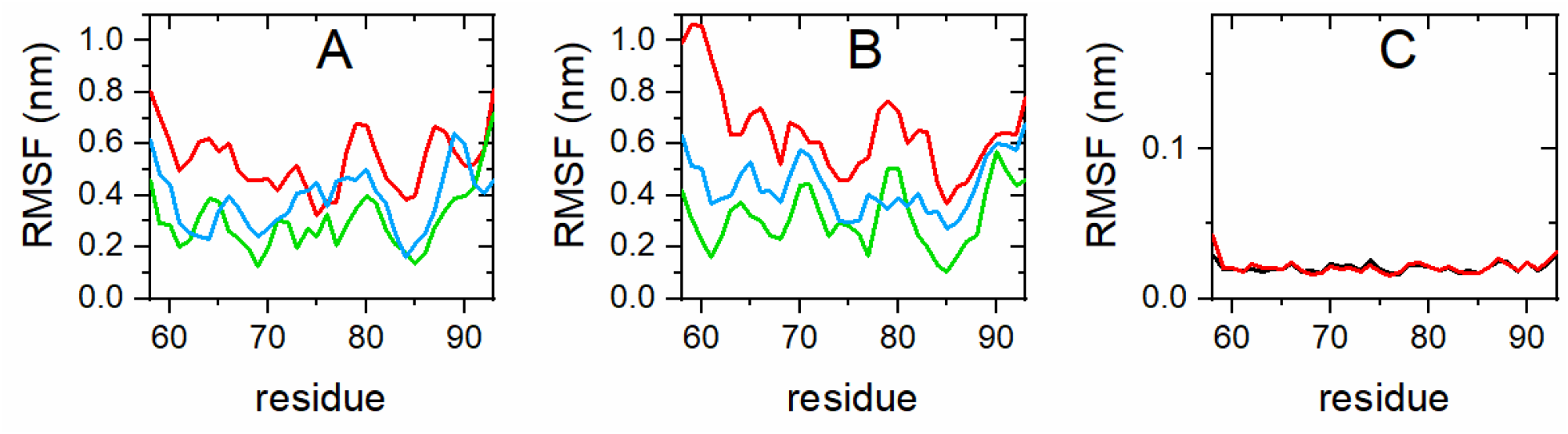
Root mean square fluctuations (RMSF) for simulations of the OR peptide Cα atoms together with: (**a**) Cu(II) ions, and (**b**) Zn(II) ions. Simulations of the OR peptide with fully protonated histidine residues and a single metal ion (i.e., either Cu(II) or Zn(II)) are shown in red, while the OR peptide with neutral histidine residues and a single metal ion are shown in green, and the OR peptide with neutral histidine residues and two metal ions are shown in blue. RMSF for (**c**) two OR peptide molecules simulated with neutral histidine residues and a single Cu(II) ion. OR-1 data is black and OR-2 data red.

**Figure S6.**
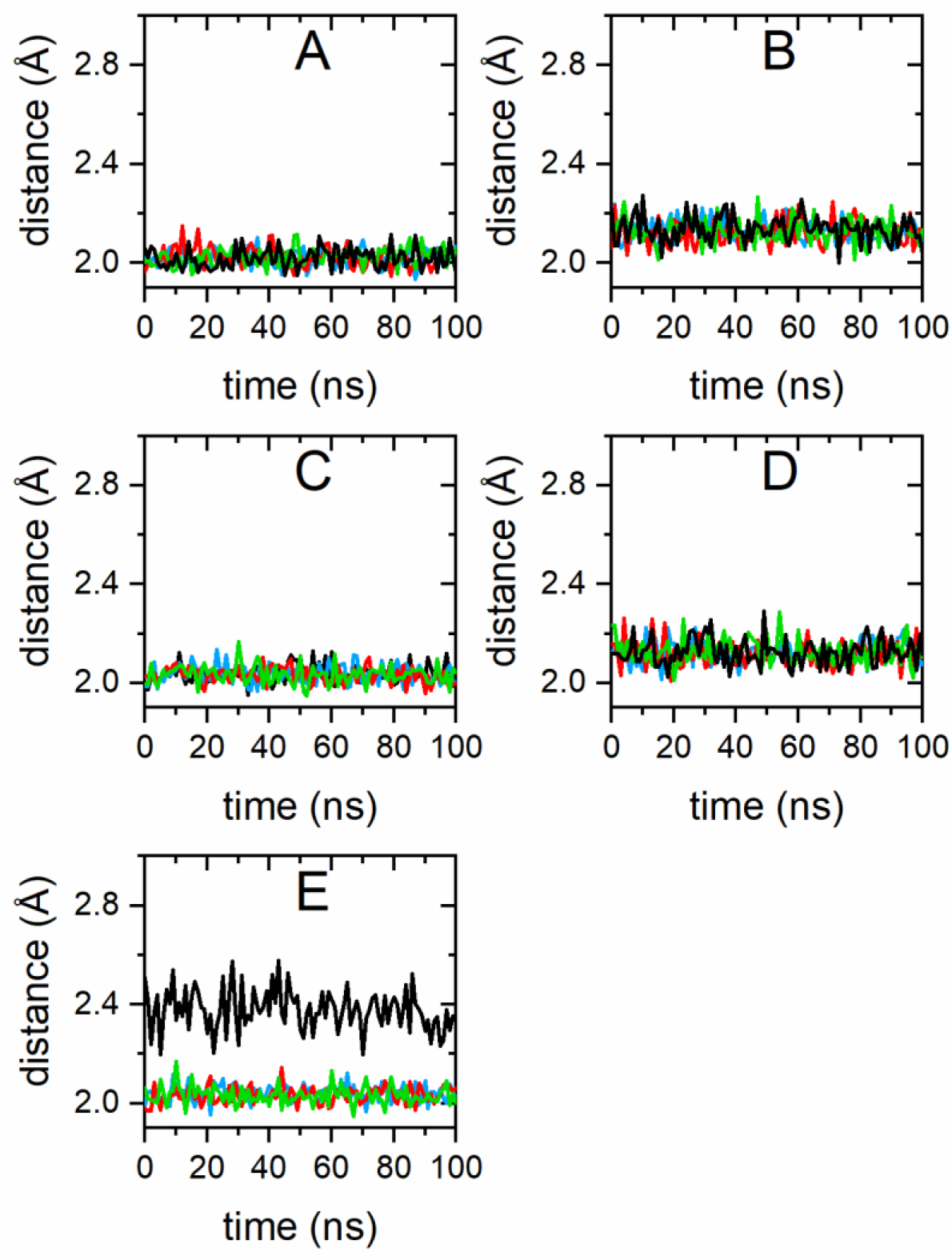
Distances between metal ions and histidine N^ε2^ atoms (H61: blue, H69: red, H77: green, H85: black) for simulations of the OR peptide together with: (**a**) a single Cu(II) ion; (**b**) two Cu(II) ions; (**c**) a single Zn(II) ion; (**d**) two Zn(II) ions, and (**e**) two OR peptide molecules with a single Cu(II) ion.

## References

1. Prusiner, S. B. Nobel Lecture: Prions. Proceedings of the National Academy of Sciences 95, 13363–13383 (1998).

2. Colby, D. W. & Prusiner, S. B. Prions. Cold Spring Harbor Perspectives in Biology 3, a006833–a006833 (2011).

3. Zahn, R. et al. NMR solution structure of the human prion protein. Proceedings of the National Academy of Sciences 97, 145–150 (2000).

4. Stahl, N. Scrapie prion protein contains a phosphatidylinositol glycolipid. Cell 51, 229–240 (1987).

5. Herms, J. et al. Evidence of Presynaptic Location and Function of the Prion Protein. J. Neurosci. 19, 8866–8875 (1999).

6. Abelein, A. et al. The hairpin conformation of the amyloid β peptide is an important structural motif along the aggregation pathway. J Biol Inorg Chem 19, 623–634 (2014).

7. Selkoe, D. J. & Hardy, J. The amyloid hypothesis of Alzheimer’s disease at 25 years. EMBO Mol Med 8, 595–608 (2016).

8. Saá, P., Harris, D. A. & Cervenakova, L. Mechanisms of prion-induced neurodegeneration. Expert Rev. Mol. Med. 18, e5 (2016).

9. Chen, C. & Dong, X.-P. Epidemiological characteristics of human prion diseases. Infect Dis Poverty 5, 47 (2016).

10. Prusiner, S. B. A Unifying Role for Prions in Neurodegenerative Diseases. Science 336, 1511–1513 (2012).

11. Collinge, J. Mammalian prions and their wider relevance in neurodegenerative diseases. Nature 539, 217–226 (2016).

12. Miller, G. Could They All Be Prion Diseases? Science 326, 1337–1339 (2009).

13. Sabate, R. When amyloids become prions. Prion 8, 233–239 (2014).

14. Walker, L. C. Prion-like mechanisms in Alzheimer disease. in Handbook of Clinical Neurology vol. 153 303–319 (Elsevier, 2018).

15. Luo, J., Wärmländer, S. K. T. S., Gräslund, A. & Abrahams, J. P. Cross-interactions between the Alzheimer Disease Amyloid-β Peptide and Other Amyloid Proteins: A Further Aspect of the Amyloid Cascade Hypothesis. Journal of Biological Chemistry 291, 16485–16493 (2016).

16. Ma, J., Gao, J., Wang, J. & Xie, A. Prion-Like Mechanisms in Parkinson’s Disease. Front. Neurosci. 13, 552 (2019).

17. Nonaka, T. & Hasegawa, M. TDP-43 Prions. Cold Spring Harb Perspect Med 8, a024463 (2018).

18. Koski, L., Ronnevi, C., Berntsson, E., Wärmländer, S. K. T. S. & Roos, P. M. Metals in ALS TDP-43 Pathology. IJMS 22, 12193 (2021).

19. Corbett, G. T. et al. PrP is a central player in toxicity mediated by soluble aggregates of neurodegeneration-causing proteins. Acta Neuropathol 139, 503–526 (2020).

20. Linden, R. et al. Physiology of the Prion Protein. Physiological Reviews 88, 673–728 (2008).

21. Schmitt-Ulms, G., Ehsani, S., Watts, J. C., Westaway, D. & Wille, H. Evolutionary Descent of Prion Genes from the ZIP Family of Metal Ion Transporters. PLoS ONE 4, e7208 (2009).

22. Toni, M., Massimino, M. L., De Mario, A., Angiulli, E. & Spisni, E. Metal Dyshomeostasis and Their Pathological Role in Prion and Prion-Like Diseases: The Basis for a Nutritional Approach. Front. Neurosci. 11, (2017).

23. Kawahara, M., Tanaka, K. & Mizuno, D. Disruption of Metal Homeostasis and the Pathogenesis of Prion Diseases. in Prion - An Overview (ed. Tutar, Y.) (InTech, 2017). doi:10.5772/67327.

24. Watt, N. T., Griffiths, H. H. & Hooper, N. M. Neuronal zinc regulation and the prion protein. Prion 7, 203–208 (2013).

25. Gielnik, M. et al. Zn(II) binding causes interdomain changes in the structure and flexibility of the human prion protein. Sci Rep 11, 21703 (2021).

26. Brown, D. R. et al. The cellular prion protein binds copper in vivo. Nature 390, 684–687 (1997).

27. Younan, N. D. et al. Copper(II)-Induced Secondary Structure Changes and Reduced Folding Stability of the Prion Protein. Journal of Molecular Biology 410, 369–382 (2011).

28. Nowakowski, M. et al. Electronic properties of a PrP ^C^–Cu(II) complex as a marker of 5-fold Cu(II) coordination. Metallomics 11, 632–642 (2019).

29. Jackson, G. S. et al. Location and properties of metal-binding sites on the human prion protein. Proceedings of the National Academy of Sciences 98, 8531–8535 (2001).

30. Remelli, M. et al. Thermodynamic and spectroscopic investigation on the role of Met residues in CuII binding to the non-octarepeat site of the human prion protein. Metallomics 4, 794 (2012).

31. Jones, C. E., Abdelraheim, S. R., Brown, D. R. & Viles, J. H. Preferential Cu2+ Coordination by His96 and His111 Induces β-Sheet Formation in the Unstructured Amyloidogenic Region of the Prion Protein. Journal of Biological Chemistry 279, 32018–32027 (2004).

32. Kardos, J. et al. Copper signalling: causes and consequences. Cell Commun Signal 16, 71 (2018).

33. Hopt, A. et al. Methods for studying synaptosomal copper release. Journal of Neuroscience Methods 128, 159–172 (2003).

34. Kardos, J., Kovács, I., Hajós, F., Kálmán, M. & Simonyi, M. Nerve endings from rat brain tissue release copper upon depolarization. A possible role in regulating neuronal excitability. Neuroscience Letters 103, 139–144 (1989).

35. Viles, J. H. et al. Copper binding to the prion protein: Structural implications of four identical cooperative binding sites. Proceedings of the National Academy of Sciences 96, 2042–2047 (1999).

36. Burns, C. S. et al. Molecular Features of the Copper Binding Sites in the Octarepeat Domain of the Prion Protein ^†^. Biochemistry 41, 3991–4001 (2002).

37. Chattopadhyay, M. et al. The Octarepeat Domain of the Prion Protein Binds Cu(II) with Three Distinct Coordination Modes at pH 7.4. J. Am. Chem. Soc. 127, 12647–12656 (2005).

38. Walter, E. D., Chattopadhyay, M. & Millhauser, G. L. The Affinity of Copper Binding to the Prion Protein Octarepeat Domain: Evidence for Negative Cooperativity ^†^. Biochemistry 45, 13083–13092 (2006).

39. Millhauser, G. L. Copper Binding in the Prion Protein ^†^. Acc. Chem. Res. 37, 79–85 (2004).

40. Millhauser, G. L. Chapter 12. Copper and Prion Protein Function: A Brief Review of Emerging Theories of Neuroprotection. in Drug Discovery (eds. Milardi, D. & Rizzarelli, E.) 249–258 (Royal Society of Chemistry, 2011). doi:10.1039/9781849733014-00249.

41. Millhauser, G. L. Copper and the Prion Protein: Methods, Structures, Function, and Disease. Annu. Rev. Phys. Chem. 58, 299–320 (2007).

42. Salzano, G., Giachin, G. & Legname, G. Structural Consequences of Copper Binding to the Prion Protein. Cells 8, 770 (2019).

43. Kambe, T., Tsuji, T., Hashimoto, A. & Itsumura, N. The Physiological, Biochemical, and Molecular Roles of Zinc Transporters in Zinc Homeostasis and Metabolism. Physiological Reviews 95, 749–784 (2015).

44. Assaf, S. Y. & Chung, S.-H. Release of endogenous Zn2+ from brain tissue during activity. Nature 308, 734–736 (1984).

45. Perera, W. S. S. & Hooper, N. M. Ablation of the metal ion-induced endocytosis of the prion protein by disease-associated mutation of the octarepeat region. Current Biology 11, 519–523 (2001).

46. Watt, N. T. et al. Prion protein facilitates uptake of zinc into neuronal cells. Nat Commun 3, 1134 (2012).

47. Walter, E. D., Stevens, D. J., Visconte, M. P. & Millhauser, G. L. The Prion Protein is a Combined Zinc and Copper Binding Protein: Zn ^2+^ Alters the Distribution of Cu ^2+^ Coordination Modes. J. Am. Chem. Soc. 129, 15440–15441 (2007).

48. Markham, K. A., Roseman, G. P., Linsley, R. B., Lee, H.-W. & Millhauser, G. L. Molecular Features of the Zn2+ Binding Site in the Prion Protein Probed by 113Cd NMR. Biophysical Journal 116, 610–620 (2019).

49. Watt, N. T. & Hooper, N. M. The prion protein and neuronal zinc homeostasis. Trends in Biochemical Sciences 28, 406–410 (2003).

50. Gielnik, M. et al. PrP (58–93) peptide from unstructured N-terminal domain of human prion protein forms amyloid-like fibrillar structures in the presence of Zn ^2+^ ions. RSC Adv. 9, 22211–22219 (2019).

51. Kelly, M. A. et al. Host-Guest Study of Left-Handed Polyproline II Helix Formation ^†^. Biochemistry 40, 14376–14383 (2001).

52. Abraham, M. J. et al. GROMACS: High performance molecular simulations through multi-level parallelism from laptops to supercomputers. SoftwareX 1–2, 19–25 (2015).

53. Kaminski, G. A., Friesner, R. A., Tirado-Rives, J. & Jorgensen, W. L. Evaluation and Reparametrization of the OPLS-AA Force Field for Proteins via Comparison with Accurate Quantum Chemical Calculations on Peptides ^†^. J. Phys. Chem. B 105, 6474–6487 (2001).

54. Jorgensen, W. L., Chandrasekhar, J., Madura, J. D., Impey, R. W. & Klein, M. L. Comparison of simple potential functions for simulating liquid water. The Journal of Chemical Physics 79, 926–935 (1983).

55. Bondi, A. van der Waals Volumes and Radii. J. Phys. Chem. 68, 441–451 (1964).

56. Hess, B., Bekker, H., Berendsen, H. J. C. & Fraaije, J. G. E. M. LINCS: A linear constraint solver for molecular simulations. Journal of Computational Chemistry 18, 1463–1472 (1997).

57. Miyamoto, S. & Kollman, P. A. Settle: An analytical version of the SHAKE and RATTLE algorithm for rigid water models. J. Comput. Chem. 13, 952–962 (1992).

58. Liao, Q., Kamerlin, S. C. L. & Strodel, B. Development and Application of a Nonbonded Cu ^2+^ Model That Includes the Jahn–Teller Effect. J. Phys. Chem. Lett. 6, 2657–2662 (2015).

59. Pogostin, B. H., Malmendal, A., Londergan, C. H. & Åkerfeldt, K. S. pKa Determination of a Histidine Residue in a Short Peptide Using Raman Spectroscopy. Molecules 24, 405 (2019).

60. Bussi, G., Donadio, D. & Parrinello, M. Canonical sampling through velocity rescaling. The Journal of Chemical Physics 126, 014101 (2007).

61. Parrinello, M. & Rahman, A. Polymorphic transitions in single crystals: A new molecular dynamics method. Journal of Applied Physics 52, 7182–7190 (1981).

62. Essmann, U. et al. A smooth particle mesh Ewald method. The Journal of Chemical Physics 103, 8577–8593 (1995).

63. Humphrey, W., Dalke, A. & Schulten, K. VMD: Visual molecular dynamics. Journal of Molecular Graphics 14, 33–38 (1996).

64. Srinivasan, R. & Rose, G. D. A physical basis for protein secondary structure. Proceedings of the National Academy of Sciences 96, 14258–14263 (1999).

65. Skjærven, L., Yao, X.-Q., Scarabelli, G. & Grant, B. J. Integrating protein structural dynamics and evolutionary analysis with Bio3D. BMC Bioinformatics 15, 399 (2014).

66. Bas, D. C., Rogers, D. M. & Jensen, J. H. Very fast prediction and rationalization of pKa values for protein-ligand complexes. Proteins 73, 765–783 (2008).

67. Sitkoff, D., Sharp, K. A. & Honig, B. Accurate Calculation of Hydration Free Energies Using Macroscopic Solvent Models. J. Phys. Chem. 98, 1978–1988 (1994).

68. Pahari, S., Sun, L., Basu, S. & Alexov, E. DelPhiPKa: Including salt in the calculations and enabling polar residues to titrate. Proteins 86, 1277–1283 (2018).

69. Case, D. A. et al. The Amber biomolecular simulation programs. J. Comput. Chem. 26, 1668–1688 (2005).

70. Smith, C. J. et al. Conformational properties of the prion octa-repeat and hydrophobic sequences. FEBS Letters 405, 378–384 (1997).

71. Jenness, D. D., Sprecher, C. & Johnson, W. C. Circular dichroism of collagen, gelatin, and poly(proline) II in the vacuum ultraviolet. Biopolymers 15, 513–521 (1976).

72. Di Natale, G. et al. Membrane Interactions and Conformational Preferences of Human and Avian Prion N-Terminal Tandem Repeats: The Role of Copper(II) Ions, pH, and Membrane Mimicking Environments. J. Phys. Chem. B 114, 13830–13838 (2010).

73. Garnett, A. P. & Viles, J. H. Copper Binding to the Octarepeats of the Prion Protein: AFFINITY, SPECIFICITY, FOLDING, AND COOPERATIVITY: INSIGHTS FROM CIRCULAR DICHROISM. J. Biol. Chem. 278, 6795–6802 (2003).

74. Taubner, L. M., Bienkiewicz, E. A., Copié, V. & Caughey, B. Structure of the Flexible Amino-Terminal Domain of Prion Protein Bound to a Sulfated Glycan. Journal of Molecular Biology 395, 475–490 (2010).

75. Danielsson, J., Jarvet, J., Damberg, P. & Gräslund, A. The Alzheimer β-peptide shows temperature-dependent transitions between left-handed 31-helix, β-strand and random coil secondary structures: Structural transitions of Alzheimer β-peptide. FEBS Journal 272, 3938–3949 (2005).

76. Ranjbar, B. & Gill, P. Circular Dichroism Techniques: Biomolecular and Nanostructural Analyses-A Review. Chemical Biology & Drug Design 74, 101–120 (2009).

77. Micsonai, A. et al. Accurate secondary structure prediction and fold recognition for circular dichroism spectroscopy. Proc Natl Acad Sci USA 112, E3095–E3103 (2015).

78. Kramer, M. L. et al. Prion Protein Binds Copper within the Physiological Concentration Range. J. Biol. Chem. 276, 16711–16719 (2001).

79. Stöckel, J., Safar, J., Wallace, A. C., Cohen, F. E. & Prusiner, S. B. Prion Protein Selectively Binds Copper(II) Ions ^†^. Biochemistry 37, 7185–7193 (1998).

80. Wells, M. A. et al. Multiple forms of copper (II) co-ordination occur throughout the disordered N-terminal region of the prion protein at pH 7.4. Biochemical Journal 400, 501–510 (2006).

81. Good, N. E. et al. Hydrogen Ion Buffers for Biological Research *. Biochemistry 5, 467–477 (1966).

82. Hansen, A. L. & Kay, L. E. Measurement of histidine pKa values and tautomer populations in invisible protein states. Proceedings of the National Academy of Sciences 111, E1705–E1712 (2014).

83. Zahn, R. The Octapeptide Repeats in Mammalian Prion Protein Constitute a pH-dependent Folding and Aggregation Site. Journal of Molecular Biology 334, 477–488 (2003).

84. Whittal, R. M. et al. Copper binding to octarepeat peptides of the prion protein monitored by mass spectrometry. Protein Science 9, 332–343 (2000).

85. Moehl, W., Schweiger, A. & Motschi, H. Modes of phosphate binding to copper(II): investigations of the electron spin echo envelope modulation of complexes on surfaces and in solutions. Inorg. Chem. 29, 1536–1543 (1990).

86. Penido, M. G. M. G. & Alon, U. S. Phosphate homeostasis and its role in bone health. Pediatr Nephrol 27, 2039–2048 (2012).

87. Spevacek, A. R. et al. Zinc Drives a Tertiary Fold in the Prion Protein with Familial Disease Mutation Sites at the Interface. Structure 21, 236–246 (2013).

88. Pushie, M. J., Rauk, A., Jirik, F. R. & Vogel, H. J. Can copper binding to the prion protein generate a misfolded form of the protein? Biometals 22, 159–175 (2009).

89. Pushie, M. J. & Vogel, H. J. Modeling by Assembly and Molecular Dynamics Simulations of the Low Cu2+ Occupancy Form of the Mammalian Prion Protein Octarepeat Region: Gaining Insight into Cu2+-Mediated β-Cleavage. Biophysical Journal 95, 5084–5091 (2008).

90. Pushie, M. J. & Vogel, H. J. Molecular Dynamics Simulations of Two Tandem Octarepeats from the Mammalian Prion Protein: Fully Cu2+-bound and Metal-Free Forms. Biophysical Journal 93, 3762–3774 (2007).

91. Bounhar, Y., Zhang, Y., Goodyer, C. G. & LeBlanc, A. Prion Protein Protects Human Neurons against Bax-mediated Apoptosis. J. Biol. Chem. 276, 39145–39149 (2001).

92. Bocharova, O. V., Breydo, L., Salnikov, V. V. & Baskakov, I. V. Copper(II) Inhibits in Vitro Conversion of Prion Protein into Amyloid Fibrils ^†^. Biochemistry 44, 6776–6787 (2005).

93. Evans, E. G. B., Pushie, M. J., Markham, K. A., Lee, H.-W. & Millhauser, G. L. Interaction between Prion Protein’s Copper-Bound Octarepeat Domain and a Charged C-Terminal Pocket Suggests a Mechanism for N-Terminal Regulation. Structure 24, 1057–1067 (2016).

